# Phage VP882 possesses quorum-sensing-driven lytic induction and stress-mediated host growth suppression mechanisms

**DOI:** 10.64898/2026.07.27.740977

**Authors:** Grace A. Beggs, Bonnie L. Bassler

## Abstract

Vibriophage VP882 launches its lytic cascade upon detection of a quorum-sensing autoinducer produced by its bacterial host. This capability enables the phage to maximize transmission by transitioning from lysogeny to lysis only at high host cell density, when abundant host cells are present to infect. Here, we show that two different pathways can be triggered upon prophage induction. When the prophage is induced via quorum sensing, Qtip, an anti-repressor, inhibits the cI lysogeny maintenance protein, driving expression of the lysis genes. The phage can also launch a cascade that causes host growth arrest. In this case, cI derepresses two genes, one encoding a DksA homolog, TraR_VP882_, and one encoding a protein that we name QisA. TraR_VP882_ and QisA form a complex that causes host growth arrest, relying on RNAP-binding by TraR_VP882_. Phage VP882 quorum sensing also activates production of a protein we call QtiQ, which inactivates the QisA-TraR_VP882_ complex, reestablishing host cell growth. Thus, in the presence of QtiQ, the phage lysis program is enacted. To our knowledge, QtiQ is the first protein inactivator of a TraR or DksA-like homolog. By inducing growth arrest, phage VP882 may enable its host to survive under stress-inducing conditions, or the phage may delay host lysis until optimal conditions are met. In both cases, the phage thereby enhances its own prospects for spread.

---

Bacteria use the cell-to-cell communication process called quorum sensing (QS) to orchestrate collective behaviors. QS relies on the production, release, and group-wide detection of signal molecules called autoinducers^1^. Vibrios, the model genera on which the QS field was founded, possess multiple QS systems consisting of receptor-autoinducer pairs. Germane to the present work is the Vibrio QS system composed of the VqmA receptor that binds the 3,5-dimethylpyrazin-2-ol (DPO) autoinducer at high cell density^2^. This system has been most well studied in *Vibrio cholerae* in which it controls biofilm formation among other traits^3^. Phages can also participate in QS and use the information they garner to transition between lysogeny and lysis^4–7^. During lysogeny, temperate phages remain dormant and are passed down through generations as prophages. When phage lysis programs are activated, virions are produced, the current host is killed, and the virions spread to new host cells. A temperate plasmid-like vibriophage called phage VP882 possesses a homolog of VqmA (VqmA_Phage_)^8^ that launches the phage VP882 lytic cascade upon binding host-produced DPO at high host cell density^4^. DPO-bound VqmA_Phage_ activates transcription of *qtip* encoding the Qtip anti-repressor (Fig. 1a)^4,9^. Qtip binds to, oligomerizes, and inactivates the cI repressor of lysis. This activity derepresses the P_R_ promoter (Fig. 1b)^8^, which enables transcription of *q*, encoding the anti-terminator that drives expression of the phage VP882 lytic genes, *gp69-gp71*^4^ (Fig 1a, b). Analogous to other well-studied phage systems, such as phage *λ*, derepression of the P_R_ promoter also occurs following self-cleavage of cI triggered by the SOS response^4,9^. Thus, in phage VP882, derepression of the P_R_ promoter can be triggered by either QS or by DNA damage.

**Figure 1.**
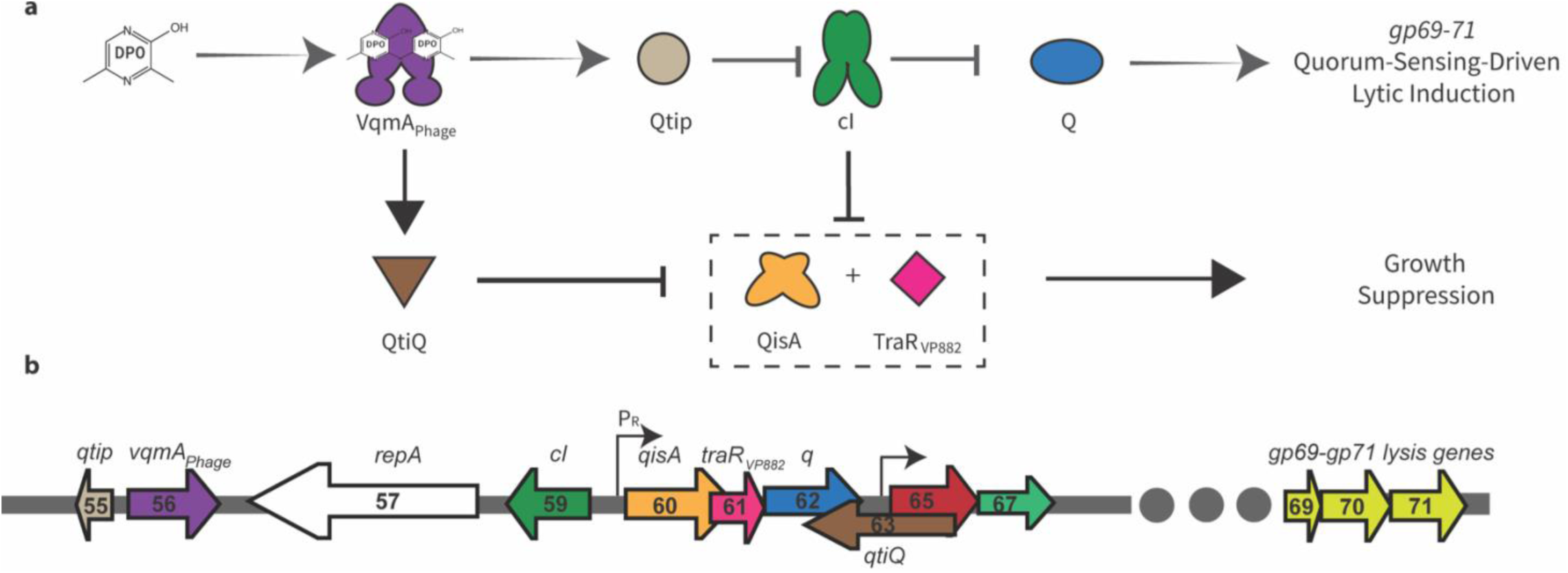
Phage VP882 possesses QS-driven lytic induction and stress-mediated host growth suppression mechanisms. **a,** Scheme showing phage VP882 QS pathway. (Top) At high host cell density, VqmA_Phage_ binds the host-produced autoinducer, DPO. The VqmA_Phage_-DPO complex activates expression of *qtip* encoding the Qtip antirepressor. Qtip inactivates the Phage VP882 cI repressor, launching the Q-mediated lytic pathway. *gp69-71* encode the lysis genes. (Bottom) The VqmA_Phage_-DPO complex activates expression of *qtiQ*, encoding QtiQ. QtiQ counteracts QisA-TraR_VP882_-mediated host growth suppression. Expression of *qisA* and *traR_VP882_* are also repressed by cI**. b,** Organization of Phage VP882 genes under study. The P_R_ promoter and the promoter driving *gp65-gp67* are denoted.

Among the phage VP882 genes under control of the P_R_ promoter is *gp61* (Fig. 1b), encoding a TraR homolog (TraR_VP882_)^4,8,10^. TraR from the *Escherichia coli* F-plasmid (TraR_F_) is a distant homolog of the stringent response protein, DksA^11^. During amino acid starvation in *E. coli* and related proteobacteria, the ppGpp alarmone accumulates and triggers the stringent response via complex formation between DksA, ppGpp, and RNA polymerase (RNAP)^12^. The consequence is increased transcription of amino acid synthesis genes and decreased transcription of genes encoding ribosomal proteins and rRNA genes. Like DksA, TraR_F_ regulates transcription by binding to the secondary channel of RNAP^11,13,14^. However, TraR_F_ does not require ppGpp for full activity^11,14^. Rather, TraR_F_ is proposed to act as a substitute for DksA to trigger the stringent response under stressful conditions in which ppGpp does not accumulate^15^. TraR proteins appear to be more widely spread than are DksA proteins as the former most frequently occur on extrachromosomal elements such as bacteriophages and plasmids, suggesting that TraR proteins could play broad roles in phage-host interactions^10^. Like TraR_F_, TraR_VP882_ activates transcription of amino acid biosynthesis genes and represses transcription of rRNA genes and genes encoding ribosomal proteins^10^. The role TraR_VP882_ plays in phage VP882 QS and how TraR_VP882_ affects phage biology have not been studied.

Here, we show that, acting synergistically with the phage Gp60 protein, TraR_VP882_ induces host growth arrest when the phage VP882 P_R_ promoter is derepressed. We name Gp60, Quorum-inhibited growth suppression assistant (QisA). We discover a companion phage VP882 protein, Gp63, that we name Quorum-triggered inactivator of QisA-TraR_VP882_ (QtiQ) and show it counteracts QisA-TraR_VP882_-mediated growth arrest. *qtiQ* is directly activated by VqmA_Phage_ and is thus under host QS control (Fig. 1a). To our knowledge, this is the first report of a protein inactivator of a TraR or a DksA-like homolog. Increased activation of phage VP882 lysis genes occurs when QtiQ is present than when absent, suggesting that phage VP882 deploys distinct mechanisms to respond to QS-signaling versus stress-signaling. Phage-driven growth arrest, by enabling infected host bacteria to evade cell death, could be a mechanism that promotes persistence of phage-infected host cells in conditions that would otherwise be lethal to the bacteria and their resident phage. Alternatively, inducing host growth arrest could be a mechanism the phage uses to delay host lysis under suboptimal conditions for virion production. In both cases, the phage could deploy these mechanisms to augment its later opportunities for spread.

## Results

### Expression of the P_R_ promoter in lysis-deficient phage VP882 induces host growth arrest

To define mechanisms underlying QS surveillance by the QS responsive phage VP882, we focused on phage processes controlled by the Qtip-cI complex. For comparison to Qtip-cI-mediated host cell lysis, we expressed *qtip* from the P*_tetA_* promoter in *V. cholerae* lysogenized with a version of phage VP882 incapable of lysis (VP882Δ*q*). Surprisingly, host growth suppression occurred (Fig. 2a). The dynamics of the reduction in host growth did not mirror those driven by induction of *qtip* in WT, i.e., in lysis-proficient phage VP882. Controls showed that Qtip is not toxic (Fig. 2a). Thus, the host growth arrest that occurred when Qtip was produced in the context of phage VP882Δ*q* must require additional phage components. Because Qtip and cI interact, we hypothesized that host growth suppression could be a downstream effect of Qtip-mediated inactivation of cI (Fig. 1a). To test this prediction, we leveraged Qtip D13A, a Qtip variant that is incapable of interacting with cI, and therefore, production of Qtip D13A does not drive expression of the cI-repressed phage VP882 P_R_ promoter (Fig 1b)^9^. Compared to WT Qtip, Qtip D13A did not cause host growth arrest in the presence of phage VP882Δ*q* (Fig. 2b and Extended Data Fig. 1a). Rather, a modest delay in lag phase occurred prior to full growth recovery. These results show that the Qtip-cI interaction is necessary to suppress host growth. We note that growth dynamics of many strains are compared throughout this work. Thus, for ease of interpretation, unlike in Fig. 2a, in Fig 2b and additional main figures, we provide data for a single time point. Full growth curves are provided in the Extended Data.

**Figure 2.**
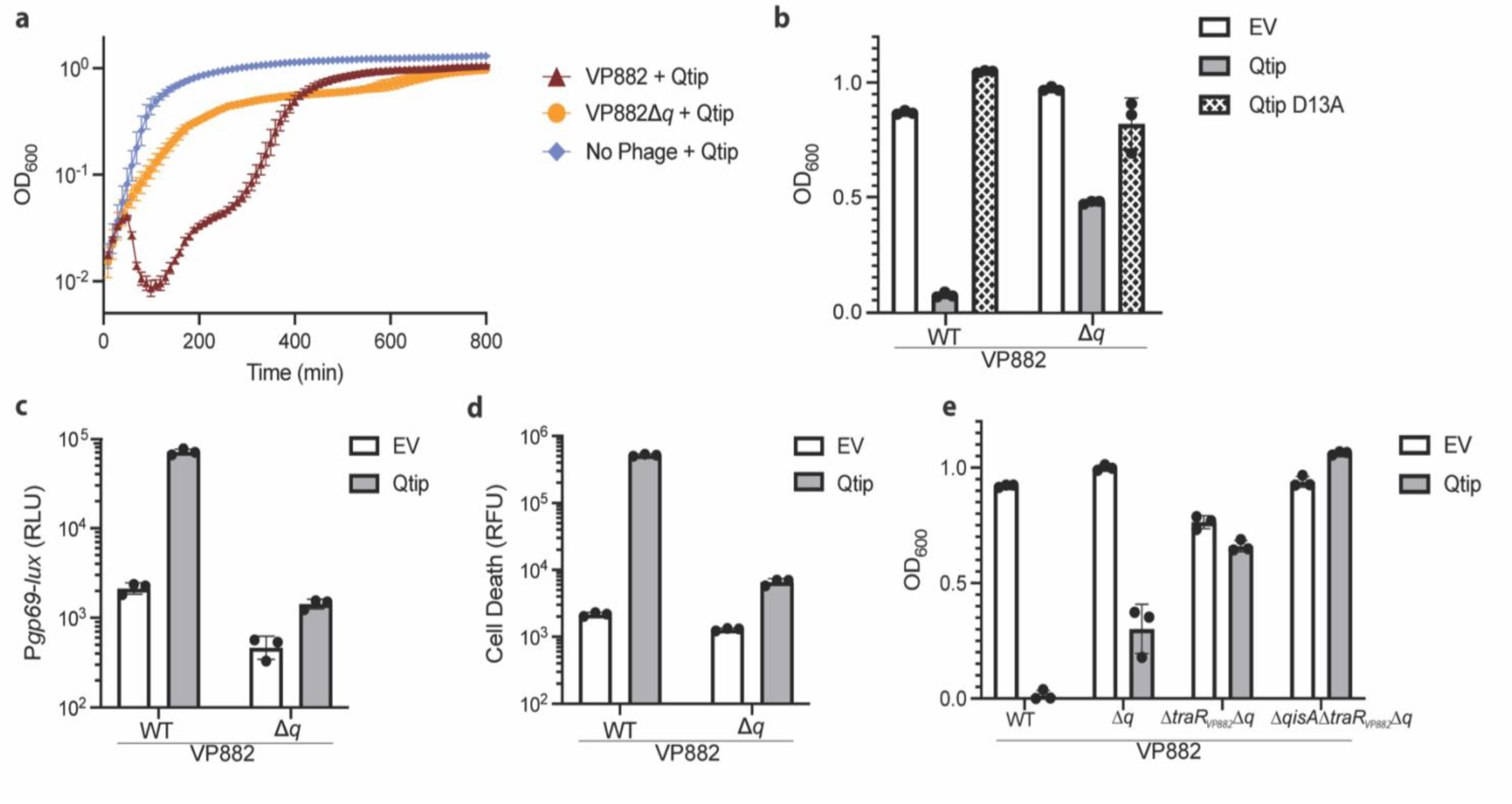
Qtip drives QisA-TraR_VP882_-mediated growth arrest in *V. cholerae* carrying lysis-deficient phage VP882. **a,** Growth curves of the designated *V. cholerae* strains carrying the indicated constructs. Qtip production was induced with 100 ng/mL aTc. **b,** OD_600_ values for *V. cholerae* lysogenized with phage VP882 or phage VP882Δ*q* carrying the designated constructs at 300 min following the addition of 100 ng/mL aTc. **c,** Bioluminescence output from P*gp69-lux* in *V. cholerae* lysogenized with phage VP882 or phage VP882Δ*q* carrying the designated constructs following addition of 200 ng/mL aTc. RLU is bioluminescence/OD_600_. **d,** Death of *V. cholerae* cells lysogenized with phage VP882 or phage VP882Δ*q* carrying the designated constructs measured by Sytox Orange fluorescence following addition of 100 ng/mL aTc. RFU is fluorescence/OD_600_. **e,** OD_600_ of *V. cholerae* carrying the designated constructs at 410 min following addition of 100 ng/mL aTc. EV denotes empty vector. Data are represented as means ± SDs with n=3 biological replicates. Full growth curves from which the values shown in panels b-e were taken are provided in Extended Data Fig. 1a-d, respectively.

The requirement for Qtip-cI interaction to limit host growth suggested that the mediator(s) of this effect could be encoded by cI-repressed phage genes. To eliminate the possibility that the downstream lysis machinery contribute to host growth arrest, we fused the luciferase (*lux*) genes to the promoter for *gp69-71*, encoding the phage VP882 cI-regulated lysis genes. Luciferase was produced following induction of *qtip* in the presence of WT phage VP882 but not phage VP882Δ*q* (Fig. 2c and Extended Data Fig. 1b). Thus, as has been shown previously^4^, Q is required for activation of *gp69-71*, but here we show that Q is dispensable for host growth arrest (Fig.2a). Thus, the phage lysis components GP69-71 cannot be responsible for restricting host cell growth.

Host growth suppression in *V. cholerae* carrying VP882Δ*q* could be a consequence of host growth inhibition or host cell death. To distinguish between these two possibilities, we quantified cell death with Sytox Orange. As expected, cell death occurred following *qtip* induction in the presence of WT phage VP882 due to activation of the *gp69-71* lytic genes. However, cell death did not occur when *qtip* was induced in the context of phage VP882Δ*q* (Fig. 2d and Extended Data Fig. 1c). Thus, Qtip-cI drives a Q-independent, lytic gene-independent, process of host growth suppression.

### Host growth suppression is mediated by QisA and TraR_VP882_

Our finding that host growth suppression depends on the Qtip-cI interaction strongly suggested that a gene or genes driven by the cI-repressed P_R_ promoter could encode the growth suppression mediator(s). Those genes are *gp60* and *gp61* and they are co-transcribed with *q*^16^ (Fig 1b). *gp60* encodes a protein of unknown function. *gp61* encodes TraR, a DksA homolog. In *E. coli*, DksA and TraR_F_, the latter from the F-plasmid, can each bind RNAP to globally alter transcription, enacting the stringent response by repressing ribosomal RNA and ribosomal protein production and activating amino acid biosynthesis genes^11,12^. A key difference between DksA and TraR_F_ is that TraR_F_ can suppress *E. coli* growth but DksA does not^14^. The mechanism underlying this difference is not known. To test whether *gp60* and/or *gp61* is involved in Qtip-cI-mediated host growth suppression, we constructed phage VP882Δ*q* lacking *gp61* or lacking both *gp60* and *gp61.* We could not generate the single VP882Δ*gp60*Δ*q* mutant because deletion of *gp60* interrupts downstream co-transcribed genes (see Fig. 1b). Introduction of these phages into *V. cholerae* revealed that the absence of *gp61* partially restored and the absence of both *gp60* and *gp61* fully restored host growth following induction of *qtip* (Fig. 2e and Extended Data Fig. 1d). Measurements of colony forming units also supported this conclusion (Extended Data Fig. 1e). From here forward we call *gp60 qisA* for quorum-inhibited growth suppression assistant; the rationale for this label will become clear below. We call *gp61 traR_VP882_*.

### QS controls production of a phage VP882 protein that counteracts QisA-TraR_VP882_-mediated growth suppression

VqmA_Phage_ activates expression of *qtip*^4^. Thus, we anticipated that inducing production of VqmA_Phage_ in the context of phage VP882Δ*q* would yield a phenotype identical to that following induction of *qtip*: host growth arrest (see Fig. 2a). Indeed, induction of *vqmA_Phage_* in *V. cholerae* carrying VP882Δ*q* or VP882Δ*traR_VP882_*Δ*q* resulted in host growth arrest and deletion of both *qisA* and *traR_VP882_* restored growth (Fig. 3a and Extended Data Fig. 2a). Curiously, however, induction of *vqmA_Phage_* in *V. cholerae* harboring phage VP882 *q*::Tn*5* did not arrest growth (Fig. 3a and Extended Data Fig. 2a), whereas induction of *qtip* in *V. cholerae* carrying phage VP882 *q*::Tn*5* did (Extended Data Fig. 2b). Thus, we hypothesized that there might be a VqmA_Phage_-activated gene encoding a function that counteracts QisA-TraR_VP882_-mediated growth suppression on phage VP882 *q*::Tn*5* but that gene is absent in phage VP882Δ*q*. Two genes fit this criterion: *gp63* and *gp64* are counter-oriented to and overlapping with the *q* gene (Fig. 3b, top). Thus, portions of both genes are deleted in phage VP882Δ*q* (Fig. 3b, bottom). Reintroduction of phage VP882 genomic fragments spanning this region into *V. cholerae* carrying phage VP882Δ*q* enabled us to pinpoint the minimal recovery region (Fig. 3b, top, designated MRR) capable of restoring growth when *vqmA_Phage_* was induced in *V. cholerae* carrying phage VP882Δ*q* (Fig. 3c and Extended Data Fig. 2c).

**Figure 3.**
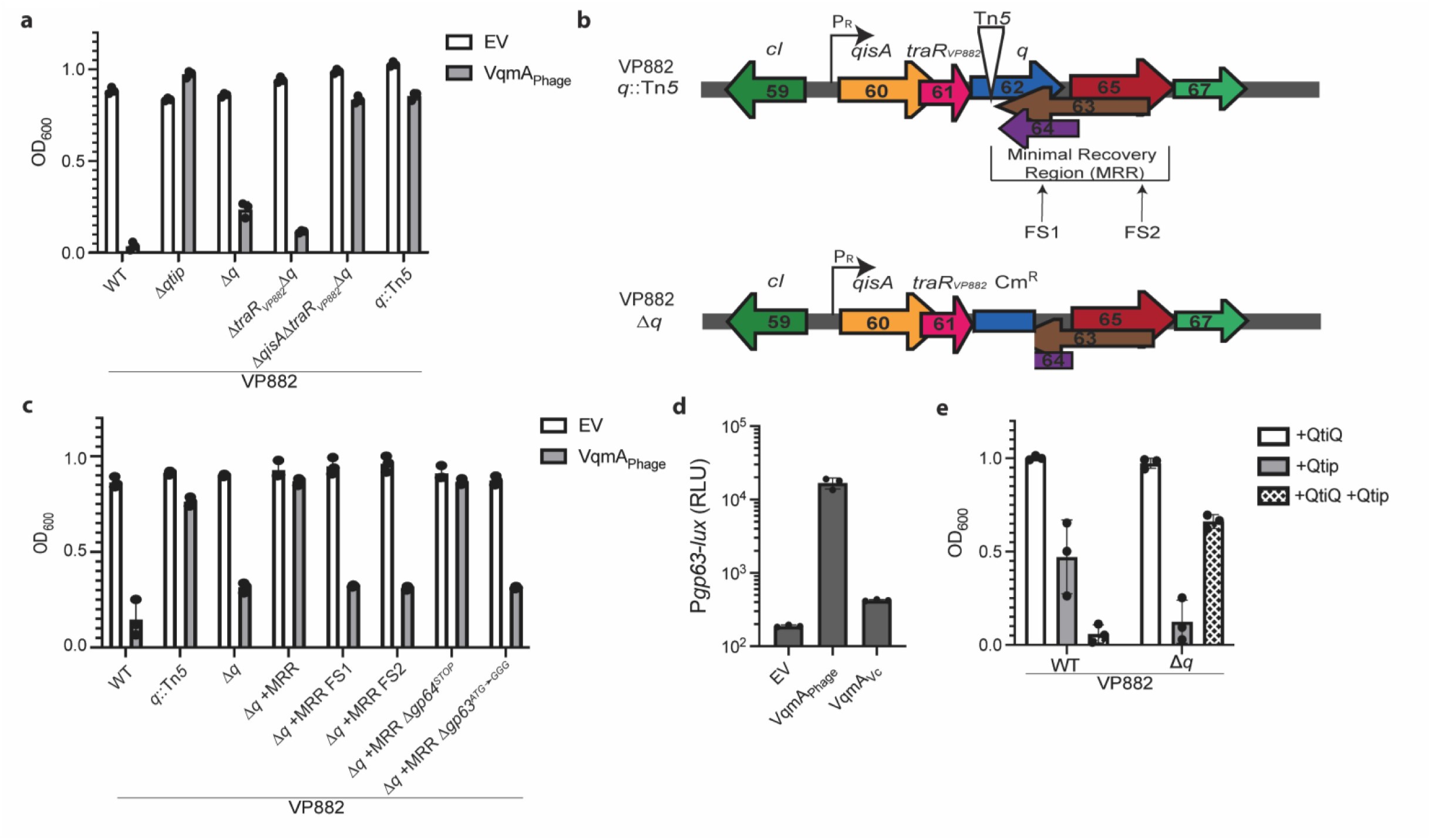
VqmA_Phage_ activates a gene encoding an inactivator of QisA-TraR_VP882_-mediated growth arrest. **a,** Growth of *V. cholerae* lysogenized with the designated phage VP882 constructs at 410 min after induction of *vqmA_Phage_*. All strains were grown with 0.2% L-arabinose. **b,** Scheme showing arrangement of genes relevant to these studies in phage VP882 *q*::Tn*5* and phage VP882Δ*q*. In phage VP882Δ*q,* two ORFs overlap the *q* gene and are disrupted by a Cm^R^ cassette. The minimal recovery region (MRR) and the two frameshift mutations (designated FS1 and FS2) tested are indicated. **c**, Growth of *V. cholerae* carrying the designated phage VP882 constructs at 410 min after induction of *vqmA_Phage_*. All strains were grown with 0.2% L-arabinose. **d**, Bioluminescence output of P*gp63-lux* in *E. coli* harboring constructs encoding the designated proteins driven by the P*_BAD_*-promoter. All strains were grown in the presence of 0.2% L-arabinose. RLU is bioluminescence/OD_600_. **e**, Growth of *V. cholerae* lysogenized with the designated versions of phage VP882 and carrying plasmids overexpressing the designated genes. *qtip* was induced with 50 ng/mL aTc. All strains were grown with 0.002% L-arabinose to induce *qtiQ* which was added following 1:5,000 back-dilution of overnight cultures. The representative time point corresponds to 600 min after *qtiQ* induction. EV designates the empty vector control. Data are represented as means ± SDs with n=3 biological replicates. Full growth curves from which the values shown in panels a, c, and e were taken are provided in Extended Data Fig. 2a, 2c, and 2f, respectively.

To establish whether the growth restoration factor encoded in the MRR is a protein or a regulatory RNA, we introduced two frameshift mutations, one into the DNA that overlaps with *q* (designated FS1) and one in the DNA downstream of *q* (designated FS2) (Fig. 3b, top). Both frameshift mutations eliminated growth recovery by the MRR (Fig. 3c and Extended Data Fig. 2c). Thus, the restoration factor is a protein. The only gene spanning both frameshift mutations is *gp63* (Fig. 3b, top). Moreover, the location of FS2 suggests that *gp63* encodes a 160-amino acid protein. To verify these assertions, we performed two tests. First, we introduced a stop codon into *gp64* that did not alter the protein sequence of Gp63. Second, we mutated the putative start codon for Gp63. Inserting a stop codon into *gp64* had no effect, while altering the Gp63 start codon eliminated growth recovery (Fig. 3c and Extended Data Fig. 2c). Our controls show that the MRR and its derivatives did not activate the lytic genes (Extended Data Fig. 2d).

To determine if VqmA_Phage_ functions directly to activate *gp63*, we constructed a P*gp63-lux* transcriptional fusion and measured bioluminescence in *E. coli* in response to production of VqmA_Phage_ (Fig. 3d). Indeed, VqmA_Phage_ drove a 100-fold increase in P*gp63-lux* output. Moreover, VqmA_Phage_ activation of P*gp63-lux* was responsive to its cognate autoinducer DPO (Extended Data Fig. 2e). Compared to VqmA_Phage_, VqmA from the host *V. cholerae* (VqmA_Vc_) did not activate P*gp63-lux* nearly as effectively suggesting that interaction with the *gp63* promoter is a rather specific property of the VqmA_Phage_ homolog (Fig. 3d). We name Gp63, QtiQ for quorum-triggered inactivator of QisA-TraR_VP882_.

Our emerging model suggests that VqmA_Phage_ directly activates *qtiQ* and QtiQ counteracts QisA-TraR_VP882_-mediated host growth suppression. If so, overexpression of *qtiQ* should bypass the necessity for VqmA_Phage_ when *qtip* is induced in *V. cholerae* carrying phage VP882Δ*q*. Fig. 3e and Extended Data Fig. 2f showed this is indeed the case: Over-production of QtiQ in this context drove nearly full growth arrest recovery. Thus, phage VP882 growth suppression is caused by QisA and TraR_VP882_ and that activity is counteracted by QtiQ, which, in turn, is controlled by QS via VqmA_Phage_-DPO (Fig. 1).

### TraR_VP882_ and QisA form a complex and the TraR_VP882_ RNAP-binding motif is dispensable for interaction with QisA but required for host growth suppression

The data in Fig. 3e suggest that QtiQ inhibits both QisA and TraR_VP882_. That, coupled with our finding that QisA and TraR_VP882_ apparently act synergistically to suppress host growth (Figs. 2e and 3a), suggested that QisA and TraR_VP882_ may form a complex. To explore this possibility, we produced HIS_6_-TraR_VP882_ and QisA-HALO in *E. coli,* used cobalt beads to pull down HIS_6_-TraR_VP882_, and evaluated the eluted fraction for QisA-HALO. Fig. 4a shows that TraR_VP882_ and QisA co-purify so they indeed form a complex. Control experiments showed that the presence of His-TraR_VP882_ was required for capture of QisA-HALO by the beads (Extended Data Fig. 3a). TraR_VP882_ has a DxxDxA motif that is highly conserved among TraR proteins and required for interaction with RNAP^10^. To determine if this motif is necessary for QisA-TraR_VP882_ complex formation, we replaced the two critical aspartate residues with asparagine residues. A complex formed between QisA-HALO and HIS_6_-TraR^NxxNxA^ (Fig. 4a) showing that the TraR_VP882_ RNAP-binding motif is dispensable for interaction with QisA.

**Figure 4.**
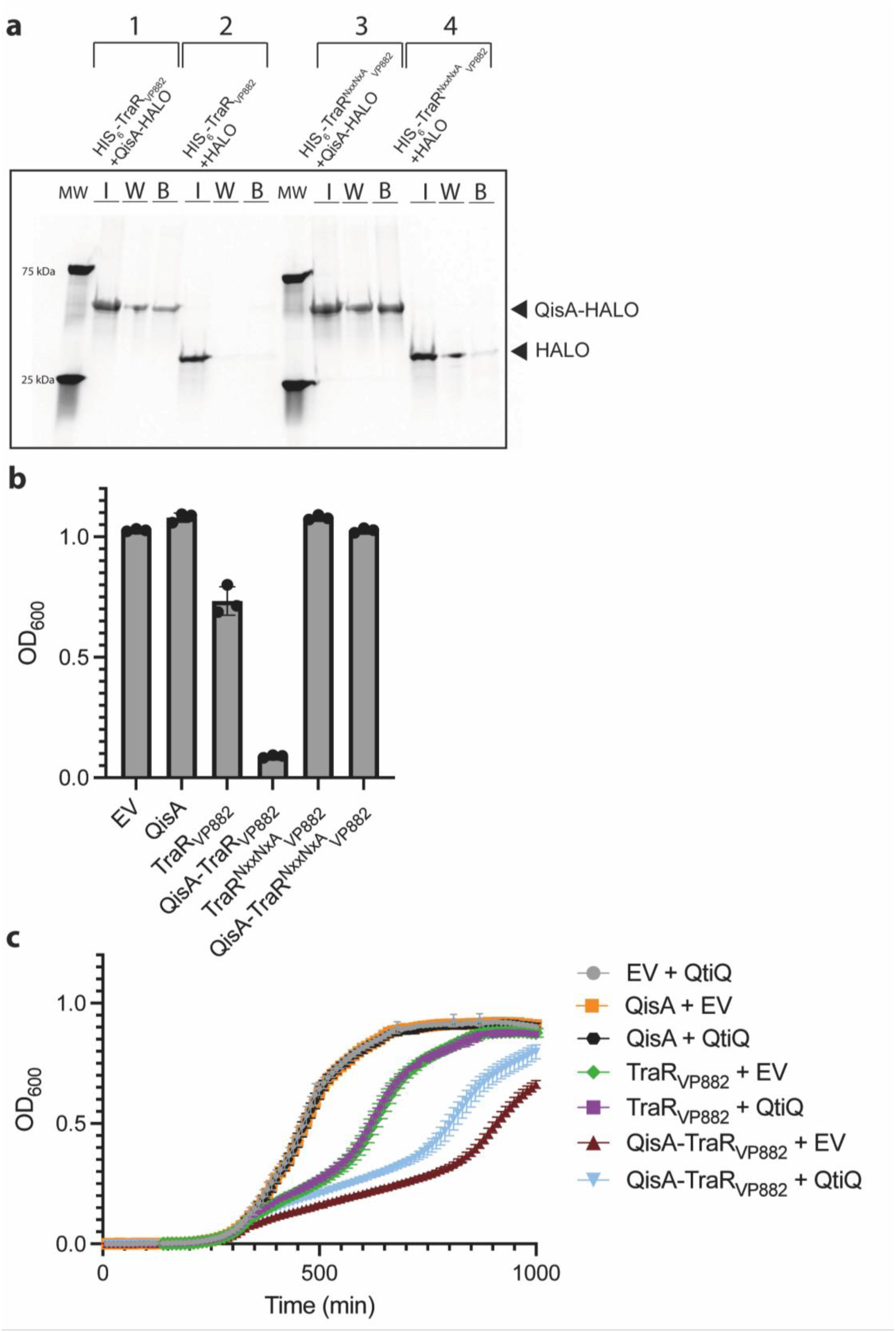
QisA and TraR_VP882_ form a complex that is sufficient to suppress host growth and for recognition by QtiQ. **a**, SDS-PAGE of proteins isolated from *E. coli* producing the following: 1, HIS_6_-TraR_VP882_ and QisA-HALO, 2, HIS_6_-TraR_VP882_ and HALO, 3, HIS_6_-TraR^NxxNxA^ and QisA-HALO, 4, HIS_6_-TraR^NxxNxA^ and HALO. I, W, and B designate input, wash, and protein retained on cobalt beads, respectively. The gel was stained for HALO. QisA-HALO and the HALO tag are shown with black arrowheads. MW designates molecular weight marker; representative bands are labeled. **b**, Growth of *E. coli* at 660 min after induction of the genes encoding the designated proteins with 50 ng/mL aTc. **c**, Growth of *E. coli* producing the designated proteins. 50 ng/mL aTc was added after 330 min. All strains were grown with 0.2% L-arabinose. EV designates empty vector. Data are represented as means ± SDs with n=3 biological replicates. Full growth curves from which values in panel b were taken are shown in Extended Data Fig. 3b.

To determine if QisA, TraR_VP882_, or the QisA-TraR_VP882_ complex is sufficient to suppress host growth, we produced them alone and together in *E. coli* (Fig. 4b and Extended Data Fig. 3b). QisA alone had no effect on growth. TraR_VP882_ caused modest growth suppression. Consistent with previous work^10^, producing other TraR homologs, i.e., TraR_λ_ or TraR_F_, in *E. coli* alone were also sufficient to yield some growth arrest (Extended Data Fig. 3c). Dramatic growth suppression occurred when TraR_VP882_ and QisA were both present (Fig. 4b and Extended Data Fig. 3b). Inducing the TraR^NxxNxA^ variant alone or with QisA eliminated growth suppression (Fig. 4b and Extended Data Fig. 3b) demonstrating that while the TraR_VP882_ RNAP-binding motif is dispensable for interaction with QisA, it is required to suppress host growth.

We were surprised that QisA did not inhibit *E. coli* growth when produced alone (Fig. 4b), unlike what occurred when TraR_VP882_ was produced alone in *E. coli* (Fig. 4b) and unlike the growth defect caused by QisA alone that occurred following induction of phage VP882Δ*traR_VP882_*Δ*q* in *V. cholerae* (Fig. 3a). This discrepancy suggested that some other factor compensates for the lack of *traR_VP882_* in *V. cholerae* carrying phage VP882Δ*traR_VP882_*Δ*q*, and together with QisA, suppresses growth. Because *traR_VP882_* was originally annotated as a conjugal transfer protein, we wondered if a conjugal component that we employ to move phage VP882 and its derivatives into *V. cholerae* was substituting for TraR_VP882_. That component would not be present in *E. coli* when we introduce QisA alone. To test this possibility, we co-produced QisA and the conjugation protein we use, TraK, which harbors a DxxDxA sequence, in *E. coli*. Indeed, growth suppression occurred (Extended Data Fig. 3c), providing a mechanistic explanation for our conundrum. Thus, while both QisA and TraR_VP882_ contribute to growth suppression, TraR_VP882_, via its RNAP-binding activity appears to be key (Fig. 4b). Additional evidence supporting this claim is provided in Extended Data Fig. 3d where we tested the individual and combined contributions of QisA, TraR_VP882_, and Q, that is, each component encoded in the cI-repressed operon, to growth suppression. Only when TraR_VP882_ was produced, either by itself or with either of the other components controlled by the P_R_ promoter, did growth suppression occur. In all cases, growth suppression was eliminated when the TraR^NxxNxA^ variant was produced instead of wild-type TraR_VP882_.

### QtiQ is sufficient to counteract QisA-TraR_VP882_-mediated growth suppression

To assess whether QtiQ is sufficient to alleviate QisA-TraR_VP882_-mediated growth suppression, we produced QtiQ with QisA, with TraR_VP882_, and with both QisA and TraR_VP882_ in *E. coli* (Fig. 4c). As shown and discussed above, producing QisA alone did not affect growth but, together, QisA and TraR_VP882_ arrested growth. QtiQ only elicited growth recovery when both QisA and TraR_VP882_ were present, not when TraR_VP882_ was produced without QisA (Fig. 4c). These data suggest that QtiQ impedes QisA-TraR_VP882_-mediated but not TraR_VP882_-mediated growth suppression.

### In the absence of QtiQ, phage VP882 driven host growth suppression rather than lysis dominates

Here, we have shown that the QisA-TraR_VP882_ complex is sufficient to induce host growth arrest and that this outcome can be circumvented by QtiQ. Curiously, VqmA_Phage_ both activates and inactivates growth suppression through activation of expression of *qtip* and *qtiQ*, respectively (Fig. 1a). Thus, to better understand the role of QtiQ in phage VP882 biology, we induced v*qmA_Phage_* in *V. cholerae* lysogenized with lysis-proficient phage VP882 in which *qtiQ* was either present (WT phage VP882) or absent (phage VP882 *qtiQ*::Tn*5*). Production of VqmA_Phage_ when WT phage VP882 was present drove the characteristic lysis curve (Fig. 5a). By contrast, growth suppression appeared to dominate when VqmA_Phage_ was produced in the context of phage VP882 *qtiQ*::Tn*5* (Fig. 5a). Indeed, as judged by the P*gp69-lux* reporter, lytic gene activation was 5-fold higher following *vqmA_Phage_* induction in the presence of WT phage VP882 compared to that when phage VP882 *qtiQ*::Tn*5* was present (Fig. 5b), suggesting that QtiQ-mediated inactivation of QisA-TraR_VP882_ promotes phage lysis. qPCR measurements of expression of the *gp69* lytic gene confirmed this result (Extended Data Fig. 4a). Comparison of levels of cell death following induction of *vqmA_Phage_* when QtiQ is present (WT phage VP882) versus absent (phage VP882 *qtiQ*::Tn*5*) also verified this assertion as there was 8.5-fold more cell death in the former (Fig. 5c). Lastly, we previously showed that phage VP882 can be induced by VqmA_Phage_ or by DNA damage.^4^ Addition of the DNA damaging compound Mitomycin C also caused growth arrest, cell death, and lytic gene activation (Extended Data Fig. 4b-d). Collectively, these results suggest that in the absence of QtiQ-driven inactivation, QisA-TraR_VP882_-mediated growth arrest dominates over phage lysis, and moreover, QtiQ is necessary to drive inactivation of QisA-TraR_VP882_-mediated growth arrest.

**Figure 5.**
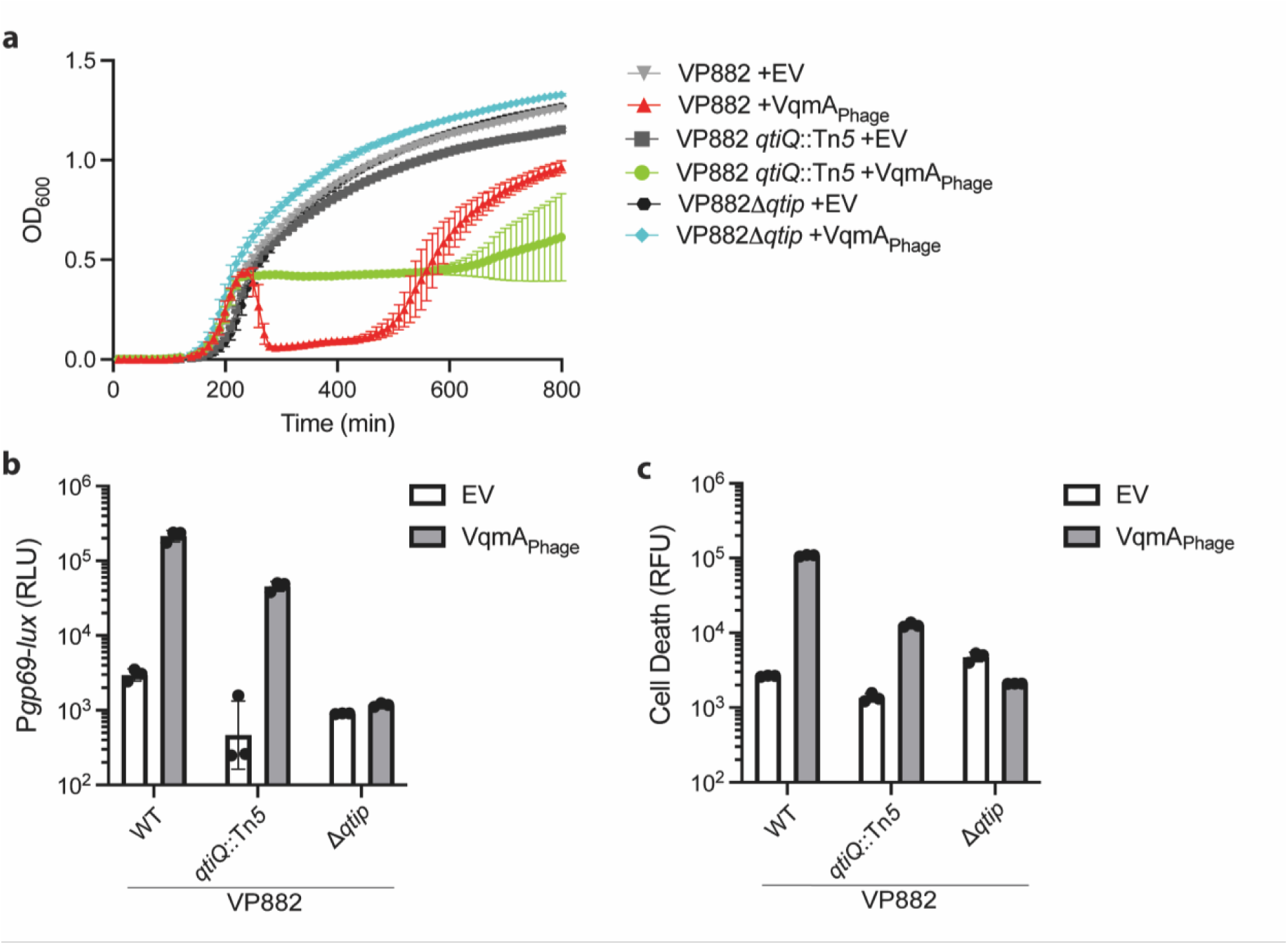
QtiQ promotes phage VP882 lysis. **a**, Growth curves of the designated *V. cholerae* strains carrying the indicated constructs. **b**, Bioluminescence output of P*gp69-lux* in *V. cholerae* lysogenized with the indicated versions of phage VP882 and carrying the designated constructs. RLU is bioluminescence/OD_600_. **c**, Cell death of *V. cholerae* lysogenized with the indicated versions of phage VP882 and carrying the designated constructs measured by Sytox Orange fluorescence. RFU is fluorescence/OD_600_. EV designates empty vector. All strains in all panels were grown in the presence of 0.002% L-arabinose to induce *vqmA_phage_*. Data are represented as means ± SDs with n=3 biological replicates.

## Discussion

We discover that the QS-responsive phage VP882 employs a host growth restriction mechanism enacted by two proteins, QisA and TraR_VP882_. We also discover that the QisA and TraR_VP882_ activities are inactivated by another QS-controlled phage protein that we name QtiQ. We show that derepression of the phage VP882 P_R_ promoter launches production of QisA and TraR_VP882_, which together form a complex that suppresses host growth via the TraR_VP882_ RNAP-binding motif. The requirement for the TraR_VP882_ RNAP-binding motif to arrest host growth suggests that QisA-TraR_VP882_ function through transcriptional control.

Previous work showed that TraR_VP882_ affects transcription of ribosomal genes and amino acid biosynthesis genes^10^. The genes affected by the QisA-TraR_VP882_ complex remain to be defined. Possibly, the QisA-TraR_VP882_ complex provides an additional layer of control over TraR_VP882_-regulated stringent response genes. We suggest this because a more severe growth defect occurs upon QisA-TraR_VP882_ production than following production of TraR_VP882_ alone (Fig. 4b). Indeed, the requirement for the conserved TraR_VP882_ DxxDxA motif to cause growth arrest suggests that the QisA-TraR_VP882_ complex must bind the RNAP secondary channel to enact control of the previously identified TraR_VP882_-regulated ribosomal and amino acid biosynthesis genes^10,13^. There may be differences between the activities of TraR_VP882_ and QisA-TraR_VP882_, analogous to the differences in the functions of *E. coli* TraR_F_ and its homolog DksA^17^. As noted above, DksA does not cause growth arrest whereas TraR proteins (e.g., TraR_F_, TraR_λ_) do^14^. Additionally, DksA promotes DNA replication^18^ and it can act as a transcription elongation factor^19^. Whether TraR proteins do likewise has not been studied. Thus, additional roles for TraR proteins and for the QisA-TraR _VP882_ complex may be revealed, and in the case of the latter, in both host and phage biology. Recently, VP882-like phages have been identified in a range of hosts and many encode *qisA* and *traR* genes^20^ suggesting the host restriction mechanism we have characterized here could be commonly employed by this phage family.

Our data show that QtiQ only inhibits QisA-TraR_VP882_-mediated growth suppression but not TraR_VP882_-mediated growth arrest. This finding suggests that QtiQ may recognize a specific binding site on QisA or on the QisA-TraR_VP882_ complex. Alternatively, QtiQ may bind RNAP to block binding of the QisA-TraR_VP882_ complex but not binding by TraR_VP882_. Inactivators of other TraR or DksA proteins may exist but have not yet been discovered because their appropriate inducing cues, QS or other stimuli, have not been present in experimental studies.

Phage-driven host growth arrest mechanisms most commonly result in either abortive infection where the host prevents the phage from completing replication because the phage triggers a host anti-phage defense system that halts growth^21,22^, or the phage blocks host DNA replication or cell division, indirectly redirecting host resources to phage replication^23–26^. Thus, growth arrest can benefit either the phage or the host. We consider how phage VP882 QisA-TraR_VP882_-mediated host growth suppression, and its inhibition by the QS-controlled inactivator QtiQ, could benefit the phage, the host, or both the phage and the host.

Regarding the phage: QisA-TraR_VP882_-mediated host growth suppression could function as a stopgap measure that slows host growth, providing more time for phage replication. Such a mechanism is analogous to that of TraR_λ_, which, by controlling transcription of host ribosomal and amino acid biosynthesis genes, indirectly increases phage burst size^10^. Phage VP882 may further fine-tune its timing of host cell lysis by subsequently deploying QtiQ to inactivate QisA-TraR_VP882_ and launch the lytic program. Indeed, under high cell density QS conditions, presumably prolonged QisA-TraR_VP882_-driven host growth suppression does not provide a reproductive benefit to the phage. To the contrary, surveilling the buildup of the DPO QS autoinducer to track increases in potential hosts in the vicinity and using that as the cue to transition from lysogeny to lysis fosters successful phage VP882 transmission.

Regarding the host: The host may recognize the QisA-TraR_VP882_ complex as an indication of phage VP882 infection, and in response, trigger an anti-phage defense system to abort infection or enter a physiological state that disfavors phage replication. To date, we have no indication that QisA-TraR_VP882_ elicits a host anti-phage defense system, and consistent with this finding, activation of phage lytic genes occurs when QisA-TraR_VP882_ are produced (Fig. 5b and Extended Data Fig. 4a). However, a recent report suggests that the *E. coli* stringent response can limit phage T7 infection^27^. To counteract this process, phage T7 deploys a portal protein that binds RelA and SpoT, the ppGpp synthesis enzymes, thereby disrupting production of the stringent-response-triggering ppGpp alarmone. Analogously, *Vibrio* hosts of phage VP882 may enter a stringent-response-like state in response to QisA-TraR_VP882_ to minimize phage replication. Phage VP882 could thwart this response by deploying QtiQ.

Regarding the host and the phage: QisA-TraR_VP882_-mediated growth suppression could be a mechanism that promotes survival of both the host and its resident prophage in response to SOS-driven cleavage of cI. Specifically, by slowing host growth and minimizing Q-mediated host cell lysis during the SOS response, lysogens carrying phage VP882 evade cell death under stress-inducing conditions. Thus, host cells survive SOS-inducing conditions via QisA-TraR_VP882_-mediated growth suppression and, moreover, because the prophage also persists, this arrangement enhances future opportunities for the phage to encounter naïve hosts and spread at high host cell density.

In conclusion, possessing distinct mechanisms to respond to QS versus SOS may maximize the ability of phage VP882 to persist and propagate. Intriguingly, these mechanisms may confer survival advantages to its host during stress. Thus, in this case, the evolutionary arms race between phages and bacteria may have yielded a relationship that, under particular conditions, is mutualistic. Other temperate phages, particularly those that integrate multiple sensory inputs from their hosts, analogous to phage VP882 detecting QS and SOS cues, may possess additional regulatory mechanisms that control processes that are mutually beneficial to the host and the phage.

## Methods

### Bacterial strains and reagents

*E. coli* and *V. cholerae* strains were grown with aeration in Luria-Bertani (LB-Miller, BD-Difco) broth at 37 °C. Strains used in this study are listed in Supplementary Table 1. Unless otherwise noted, antibiotics were used at the following concentrations: 100 µg/mL carbenicillin (Crb, Teknova), 50 µg/mL kanamycin (Kan, GoldBio), 50 µg/mL polymyxin B (Sigma), and 5 µg/mL chloramphenicol (Cm, Sigma). L-arabinose (Sigma), anhydrotetracycline (aTc, Takara Bio), 3,5-dimethylpyrazin-2-ol (DPO, AAblocks), Mitomycin C (MMC, Sigma) and isopropyl *β*-D-1-thiogalactopyranoside (IPTG, GoldBio) were used as indicated.

### Cloning Techniques

All primers and gene blocks used for plasmid construction are listed in Supplementary Table 2 and were obtained from Integrated DNA Technologies. FastCloning^28^ was used for plasmid assembly. Briefly, PCR with KOD Xtreme Hot Start DNA polymerase (Sigma) was used to generate plasmid backbone DNA and inserts. Cloning inserts and linearized plasmid backbones were combined in equimolar amounts, treated with DpnI (NEB), and added to chemically competent *E. coli* cells for plasmid assembly. All plasmids were verified by Sanger Sequencing. Plasmids are listed in Supplementary Table 3. Transfer of plasmids and recombinant phages into *V. cholerae* strains was carried out by conjugation with *E. coli* S17 followed by selective plating on LM plates supplemented with appropriate antibiotics.

### Generation of recombinant phage VP882

DNA encoding desired mutations, a Cm resistance cassette, conjugation machinery genes flanked by 3 kb of homology upstream and downstream of the target mutation to be introduced into phage VP882 were generated by splicing-by-overlap extension (SOE) PCR^29^ with iProof polymerase (Bio-Rad). Mutations were introduced into phage VP882 using natural transformation of *Vibrio parahaemolyticus* 882 using a variation of a previously described method^30^. Briefly, overnight cultures of *V. parahaemolyticus* 882 harboring pMMB*sacBtfoX* were back-diluted 1:100 in 2X Instant Ocean with 100 µM IPTG included to drive overexpression of the *V. cholerae tfoX* gene. Cultures were incubated without shaking at 30 °C for 5 h. At that point, 0.5-1 µg of PCR construct was added, and the cultures were incubated overnight at 30 °C without shaking. The next day, 1 mL of MLB medium was added to each culture, followed by incubation at 30 °C with shaking for 2 h. The cultures were plated on MMM plates with antibiotics. Isolated single colonies were selected for verification by colony PCR and Sanger sequencing. Phage VP882 carrying the desired mutations was isolated using the QIAprep Spin Miniprep Kit (Qiagen) and transferred into chemically competent *E. coli* cells by transformation. Transformants were purified by patching single colonies, verified by colony PCR and Sanger sequencing, and used for conjugating recombinant phage VP882 into *V. cholerae* as noted above.

### Growth and Bioluminescence Assays

Unless otherwise noted, overnight cultures were back-diluted 1:1,000 into fresh LB medium with appropriate antibiotics and dispensed (200 µL) into 96-well white-walled/clear bottom plates (Corning Costar). In the case of the experiment in Extended Data Fig. 2e, samples were back-diluted into M9 supplemented with 0.4% casamino acids (BD-Difco), appropriate antibiotics, and for the samples indicated, 100 µM DPO. An equal volume of water was added to samples that did not receive DPO. To induce *vqmA_Phage_* expression, L-arabinose was included in the medium upon back-dilution at the concentrations indicated in the figure legends. To induce *qtip*, cells were grown in plates at 37 °C for 90-100 min prior to addition of aTc except in the following cases: In experiments in Fig. 2e, Extended Data Fig. 1d, and Extended Data Fig. 2b, aTc was added immediately upon back-dilution. In the experiment in Fig. 3e and Extended Data Fig. 2f, cells were diluted 1:5,000 and grown in plates at 37 °C for 170 min before aTc addition. To acquire the data shown in Extended Data Fig. 4d, MMC was added after back-diluted cells were grown at 37 °C in plates for 270 min. To induce expression from P*_tetA_* in *E. coli* TOP10 for the experiments in Fig. 4b and Extended Data Fig. 3b-d, cells were grown in plates at 37 °C for 120 min prior to addition of aTc. Regarding the experiment in Fig. 4c, overnight cultures were back-diluted 1:10,000. To induce *qtiQ* expression, L-arabinose was included in the medium upon back-dilution of *E. coli* TOP10. The cells were grown in plates at 37 °C for 330 min prior to addition of aTc. In all cases, aTc, MMC, or L-arabinose was added at the concentrations indicated in the figure legends. The plates were shaken at 37 °C and optical density, or optical density and bioluminescence were measured every 10 min using a BioTek Synergy Multi-Mode plate reader. Measurement times represented in bar graphs are provided in Supplementary Table 4. Time points for growth data shown in bar graphs were selected to best represent the overall trend. Full growth curves are provided in the Extended Data. Time points for bioluminescence data were selected based on peak intensities.

### Cell Death Assay

Overnight cultures were back-diluted 1:1,000 into M9 medium supplemented with 0.4% casamino acids, appropriate antibiotics, and 500 nM Sytox Orange (Invitrogen). Aliquots (200 µL) were dispensed into 96-well black-walled/clear bottom plates (Corning Costar). In the case of *qtip* induction, cells were grown at 37 °C for 2.5 h before addition of aTc. In the case of *vqmA_Phage_* induction, L-arabinose was added at the time of back-dilution. Cells were grown at 37 °C for 315 min before MMC was added. MMC, aTc, and L-arabinose concentrations are provided in the figure legends. Plates were shaken at 37 °C and optical density and fluorescence were measured every 15 min using a BioTek Synergy Neo2 Multi-Mode plate reader. Fluorescence readings were made by excitation at 540 nm and emission at 590 nm. Measurement times represented in bar graphs are provided in Supplementary Table 4.

### Measurement of Colony Forming Units (CFU)

Overnight cultures were back-diluted 1:1,000 into fresh LB medium with appropriate antibiotics and dispensed (200 µL) into 96-well white-walled/clear bottom plates (Corning Costar). To induce *qtip*, cells were grown in plates at 37 °C for 90-100 min prior to addition of aTc. The plates were shaken at 37 °C, and after 200 min, the samples were serially diluted 1:10 – 1:100,000. 5 µL volumes of the dilutions were plated on LB plates containing Cm and Kan and grown at 37 °C overnight. Colonies were counted the next morning to calculate the CFU/mL. Images of plates were captured with an IQ800 imager (Cytiva).

### Co-Immunoprecipitation

Cultures of *E. coli* BL21(DE3) producing QisA-HALO, HIS_6_-TraR_VP882_ and QisA-HALO, HIS_6_-TraR_VP882_ and HALO, HIS_6_-TraR^NxxNxA^ and QisA-HALO, and HIS_6_-TraR^NxxNxA^ and HALO were grown overnight, back-diluted 1:50 into 50 mL of LB with Kan and grown to OD_600_ ∼0.4-0.8. At that time, 1 mM IPTG was added followed by 18 °C incubation overnight. The cells in each culture were collected by centrifugation at 4,000 RPM for 10 min at 4 °C. Cell pellets were resuspended in lysis buffer (20 mM Tris-HCl pH 7.5, 150 mM NaCl, cOmplete^TM^ protease inhibitor cocktail (Sigma)) followed by sonication. Lysates were added to 50 µL of magnetic cobalt beads (Thermo, Dynabeads) prepared in wash buffer (20 mM Tris-HCl pH 7.5, 150 mM NaCl) and incubated at room temperature for 20 min with gentle agitation. Prior to adding to the cobalt beads, total input protein concentration was determined by Bradford reagent (Bio-Rad) and all sample protein concentrations were equalized. Samples were placed on a magnetic stand separator (Thermo) to maintain proteins bound to the beads. The unbound fraction was removed with three washes of 100 µL of wash buffer followed by isolation with the magnetic stand separator. Washed beads were resuspended in 100 µL of wash buffer. Aliquots of 30 µL of the input, the first wash, and the final bead resuspension were collected and incubated with 5 µM of HALO-TMR ligand (Promega) for 20 min at room temperature, protected from light. Each aliquot was diluted with 30 µL of 4x Laemmli buffer (Bio-Rad) and incubated at 70 °C for 15 min to release protein from the beads. Samples were subjected to SDS-PAGE and imaged with an IQ800 imager (Cytiva) under the Cy3 setting.

### RT-qPCR

Overnight cultures were back-diluted 1:1,000 into medium containing 0.002% L-arabinose to induce *vqmA_Phage_.* Cultures were grown for 3.5 h, at which point the cells were collected and treated with RNAProtect Bacteria Reagent (Qiagen) according to the supplier’s protocol. Total RNA was isolated from cultures using the RNeasy Plus Mini Kit (Qiagen). RNA samples were treated with TURBO DNA-free (Invitrogen Ambion). cDNA was prepared using SuperScript IV VILO Master Mix (Invitrogen) according to the supplier’s protocol. SYBR Green mix (Quanta) and a CFX Opus 384 (Bio-Rad) were used for real-time PCR. Each cDNA sample was amplified in technical triplicate, and data were analyzed by a comparative CT method, in which the targeted lysis gene (*gp69*) was initially normalized to an internal bacterial housekeeping gene (*hfq*). Subsequently, the *gp69* fold-change for each strain was calculated by comparing the *vqmA_Phage_* data to that of the empty vector control.

### Quantitation and statistical analyses

The following software were used to collect and analyze data generated in this study: GraphPad Prism 10 for visualization and analyses of growth, luciferase, qPCR and fluorescence-based data; Gen5 for collection of growth, luciferase, and fluorescence-based data; CFX Maestro v5 for collection of RT-qPCR data; SnapGene v7 for primer design and sequence verification; and Amersham ImageQuant for imaging of SDS-PAGE gels. Data are presented as the means ± standard deviations with n = 3 biological replicates started from separate bacterial colonies measured on the same day.

## Data Availability

No sequencing or custom code is included in this paper. All materials associated with this study are deposited on Zenodo (DOI: 10.5281/zenodo.24321501). Time points corresponding to data in bar graphs are in Supplementary Table 4. Source Data for all figures are provided with this paper.

## Acknowledgements

We are grateful to members of the Bassler laboratory for insightful discussions. We thank Dr. Julie Valastyan for help with the RT-qPCR analyses.

## Funding

We acknowledge funding by the Howard Hughes Medical Institute, the National Science Foundation (grant MCB-2508324 to B.L.B.), and the National Institute of General Medical Sciences of the National Institutes of Health (F32GM149034 to G.A.B.).

The funders had no role in study design, data collection and interpretation, or preparation of the manuscript. Any opinions, findings, and conclusions or recommendations expressed in this material are those of the authors and do not necessarily reflect the views of the National Science Foundation or the National Institutes of Health. The content is solely the responsibility of the authors.

## Author Information

### Department of Molecular Biology, Princeton University, Princeton, New Jersey, USA

Grace A. Beggs & Bonnie L. Bassler

### Howard Hughes Medical Institute, Chevy Chase, Maryland, USA

Bonnie L. Bassler

### Contributions

G.A.B. and B.L.B. conceptualized the project. G.A.B. performed all experiments. G.A.B. and B.L.B. analyzed the data. G.A.B. and B.L.B. wrote the paper.

## Ethics Declarations

The authors declare no competing interests.

---

**Extended Data Fig. 1.**
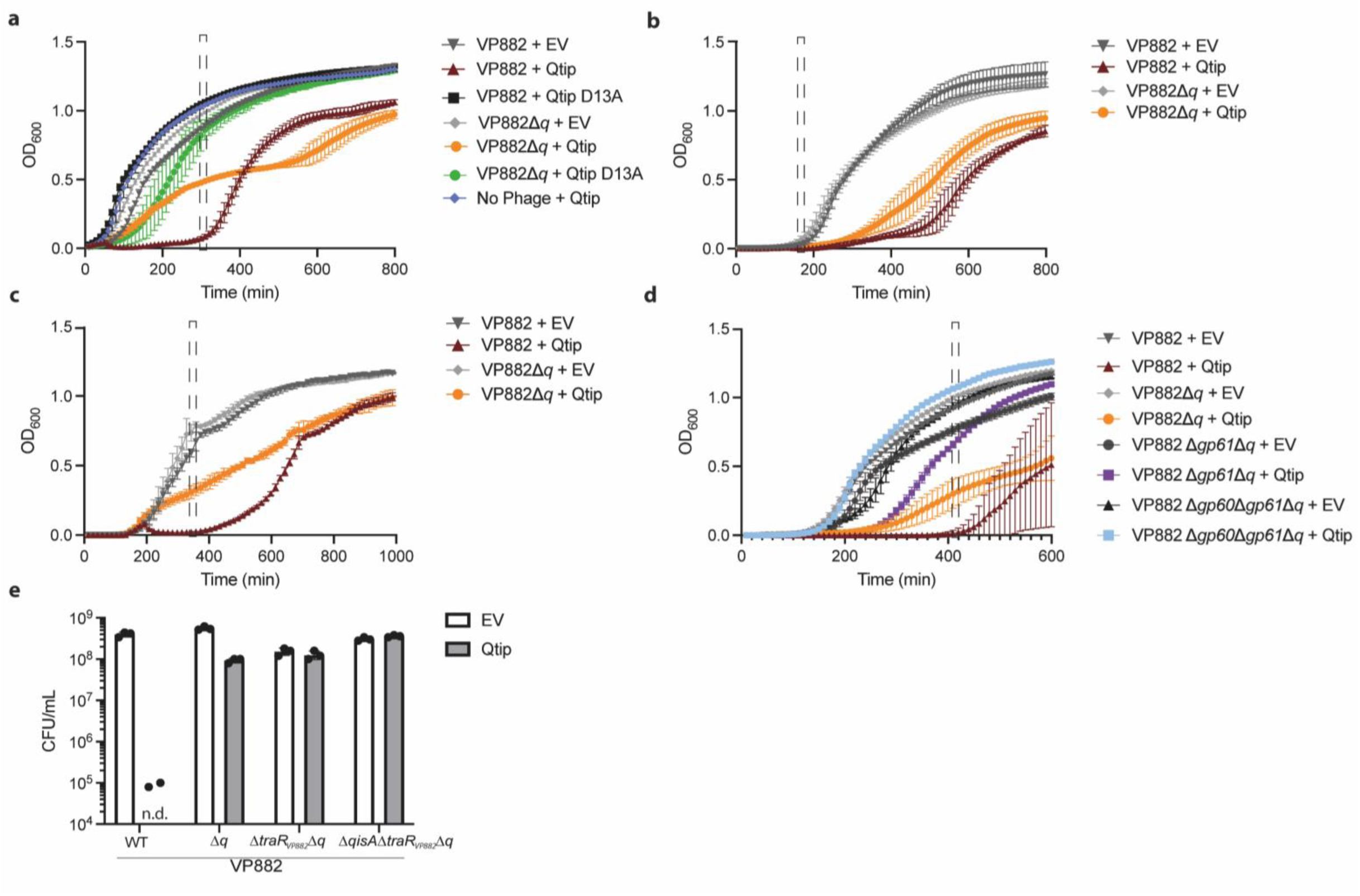
Qtip-induced host growth arrest requires interaction between Qtip and cI but does not require the phage VP882 lysis machinery. a-d, The panels show growth curves of the designated *V. cholerae* strains carrying the indicated constructs. **a**, Growth curves for which OD_600_ values are shown in Fig. 2b. **b**, Growth curves for which bioluminescence values are shown in Fig. 2c. **c**, Growth curves for which fluorescence values are shown in Fig. 2d. **d**, Growth curves for which OD_600_ values are shown in Fig. 2e. **e**, Measured colony forming units (CFU) per mL of the designated *V. cholerae* strains carrying the indicated constructs. The cultures were serially diluted and plated 200 min after the addition of 100 ng/mL aTc as described in the methods. n.d. denotes not detected. EV designates empty vector. Data are represented as means ± SDs with n=3 biological replicates. Dashed boxes in panels a-d indicate the timepoints represented in the bar graphs in Fig. 2.

**Extended Data Fig. 2.**
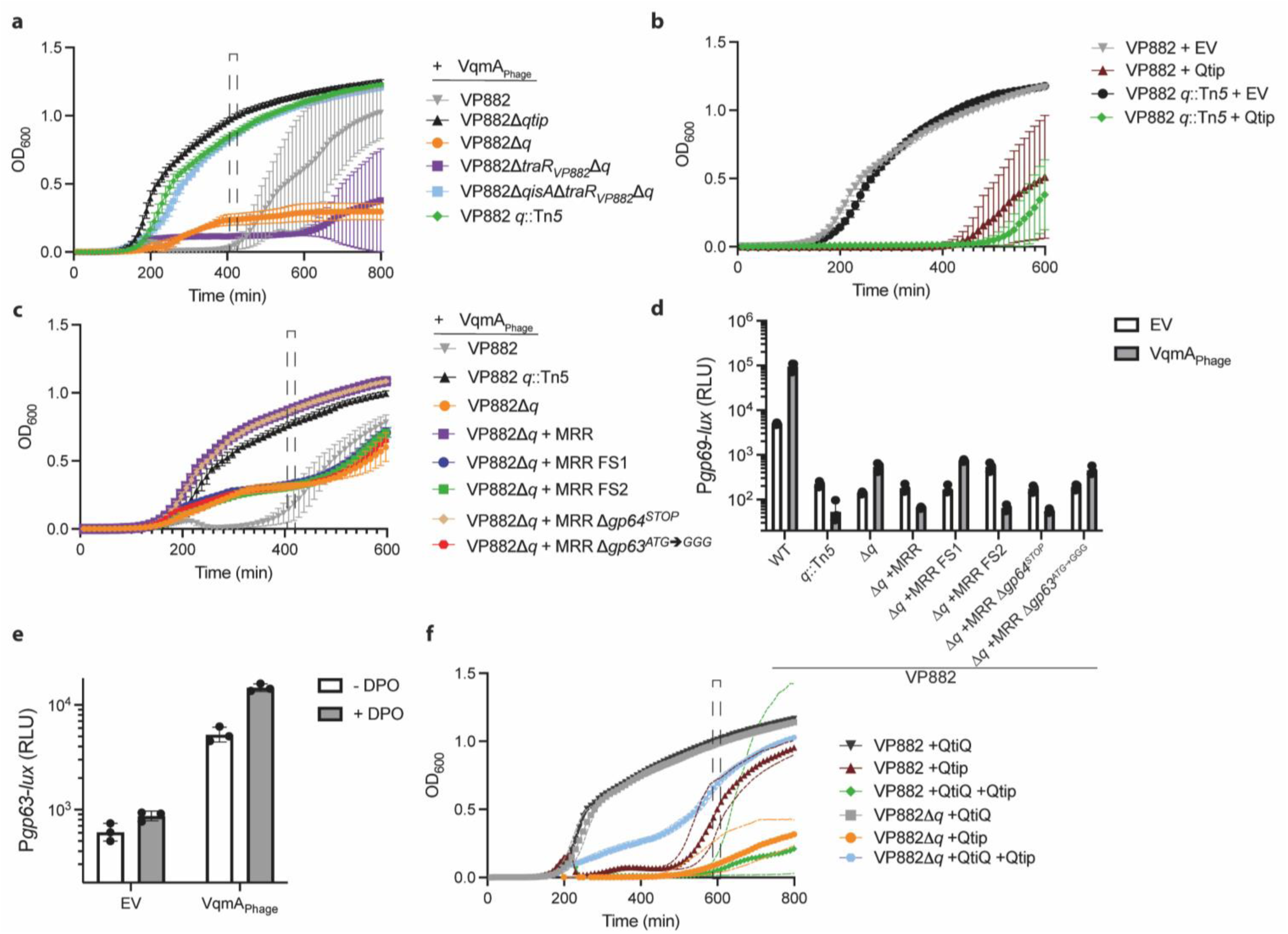
VqmA_Phage_ activates transcription of *qtiQ*, encoding a protein inhibitor of QisA-TraR_VP882_ activity. **a**, Growth curves for which OD_600_ values are shown in Fig. 3a. **b**, Growth curves of the designated *V. cholerae* strains carrying the indicated constructs treated with 100 ng/mL aTc. **c**, Growth curves for which OD_600_ values are shown in Fig. 3c. **d**, Bioluminescence output of P*gp69-lux* in the designated *V. cholerae* strains. RLU is bioluminescence/OD_600_. All strains grown with 0.2% L-arabinose. **e**, Bioluminescence output of P*gp63-lux* in Δ*tdh V. cholerae* carrying the designated constructs and grown with or without 100 µM DPO. RLU is bioluminescence/OD_600_. Tdh (threonine dehydrogenase) is required for DPO biosynthesis. All strains grown with 0.02% L-arabinose. **f**, Growth curves of the *V. cholerae* strains for which OD_600_ values are shown in Fig. 3e. In this graph, rather than error bars, we use dashed lines to show the errors. That is because providing error bars obscured the data. Solid lines show the means. Colors of dashed lines correspond to the respective solid lines. EV designates empty vector. Data are represented as means ± SDs with n=3 biological replicates. Dashed boxes in panels a, c, and f indicate the timepoints represented in Fig. 3a, 3c, and 3e, respectively.

**Extended Data Fig. 3.**
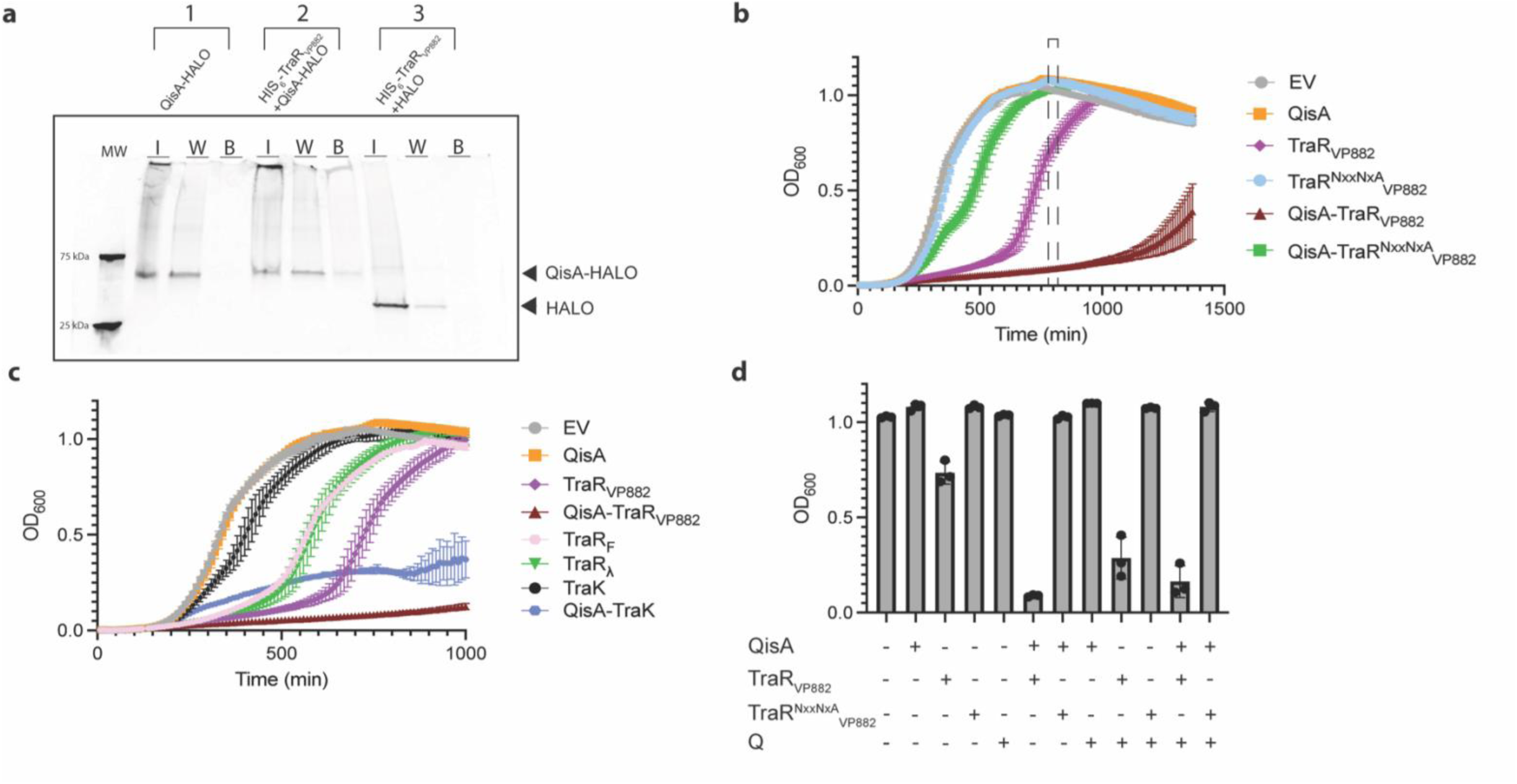
Growth suppression by QisA-TraR_VP882_ requires the TraR_VP882_ RNAP-binding motif. **a**, SDS-PAGE of proteins isolated from *E. coli* producing the following: 1, QisA-HALO alone, 2, HIS_6_-TraR_VP882_ and QisA-HALO, and 3, HIS_6_-TraR_VP882_ and HALO. I, W, and B designate input, wash, and protein retained on cobalt beads, respectively. The gel was stained for HALO. QisA-HALO and the HALO tag are shown with black arrowheads. MW designates molecular weight marker with selected bands labeled. **b**, Growth curves for which OD_600_ values are shown in Fig. 4b. The dashed box indicates the timepoint represented in Fig. 4b. **c**, Growth curves of *E. coli* carrying the genes encoding the designated proteins, which were induced with 50 ng/mL aTc at 120 min. **d**, OD_600_ values for *E. coli* at 660 min following induction of the genes encoding the designated proteins with 50 ng/mL aTc. The + and – symbols denote, respectively, the presence and absence of the gene encoding the designated protein under the control of the P*_tetA_* promoter. EV designates empty vector. Data are represented as means ± SDs with n=3 biological replicates.

**Extended Data Fig. 4.**
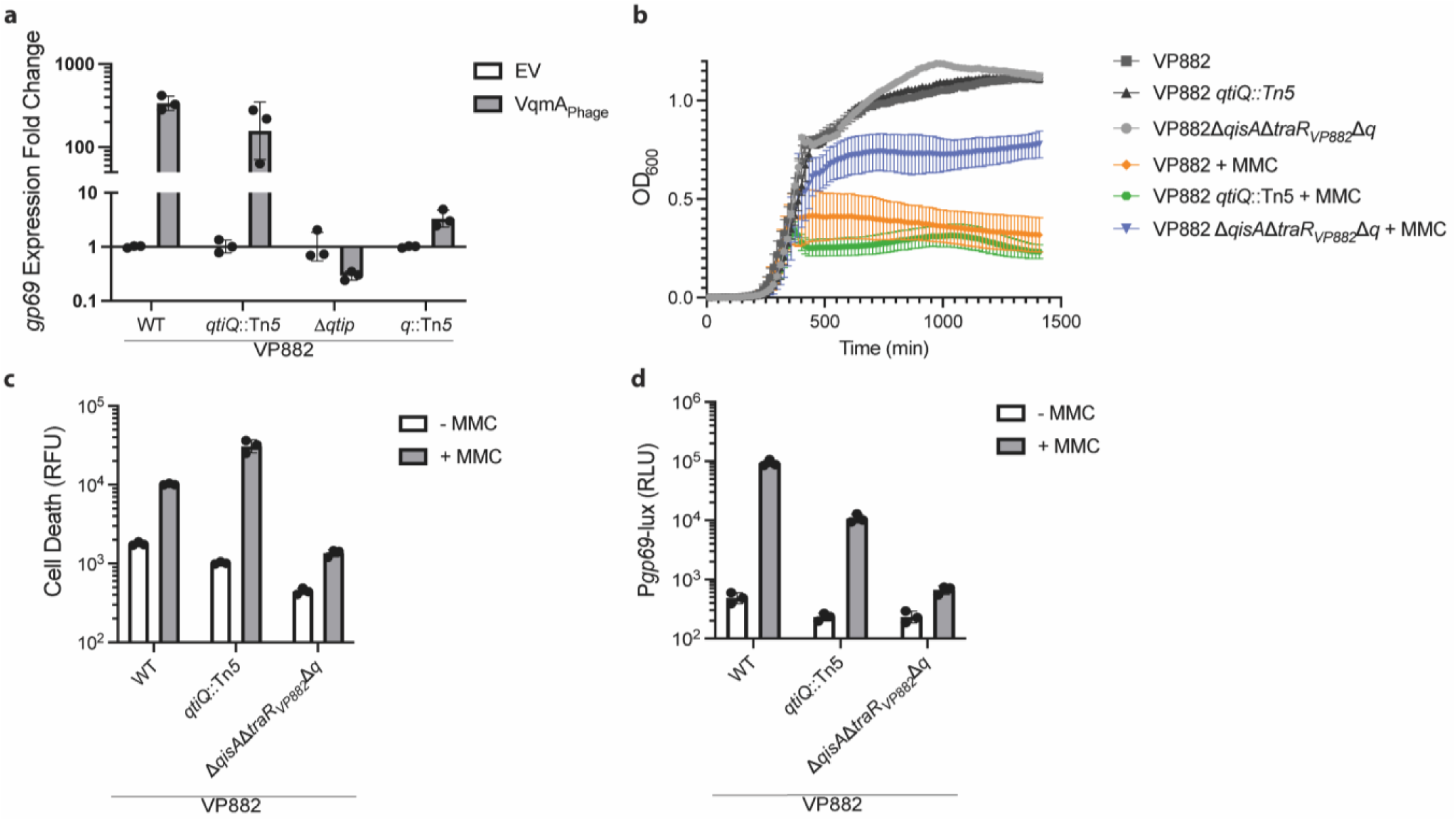
QtiQ promotes phage VP882 lytic gene activation. **a**, Fold change in *gp69* expression in *V. cholerae* strains carrying the designated phage constructs 3.5 h after induction of *vqmA_Phage_* compared to the empty vector (EV) control for each strain. Expression of *gp69* was normalized to *hfq* expression. **b**, Growth curves of *V. cholerae* carrying the designated constructs. MMC or vehicle control was added after 315 min. **c**, Cell death of *V. cholerae* lysogenized with the indicated versions of phage VP882 and carrying the designated constructs measured by Sytox Orange fluorescence 145 min after the addition of MMC. RFU is fluorescence/OD_600_. **d**, Bioluminescence output of P*gp69-lux* in *V. cholerae* lysogenized with the indicated versions of phage VP882 and carrying the designated constructs. RLU is bioluminescence/OD_600_. All strains in all panels were grown in the presence of 0.002% L-arabinose. Data are represented as means ± SDs with n=3 biological replicates. b-d, MMC was added at 50 ng/mL.

## Notes

### Competing Interest Statement

The authors have declared no competing interest.

### Summary of Updates

This version of the manuscript includes new experiments in the Extended Data Figures.

