## Supplemental Tables 1-4 for "Phage VP882 possesses quorum-sensing-driven lytic induction and stress-mediated host growth suppression mechanisms"

### Supplementary Table 1: Strains used in this study

| Parent strains | | |
|---|---|---|
| **Strain (ID)** | **Genotype** | **Reference** |
| *V. cholerae* | Wild-type C6706 | [1] |
| *V. cholerae*Δ*tdh* (KPS-842) | str. C6706, Δ*tdh* | [2] |
| *V. cholerae* (GABVc154) | str. C6706, phage VP882 *q*::Tn*5* (JSP-003) | [2] |
| *V. cholerae* (GABVc100) | str. C6706, phage VP882 *qtiQ*::Tn*5* (JSP-002) | [2] |
| *V. cholerae* (GABVc91) | str. C6706, phage VP882 $Cm^R$ (WT phage VP882) | [3] |
| *V. cholerae* (AJ030) | str. C6706, phage VP882 Δ*qtip*::$Cm^R$ | [3] |
| *V. cholerae* (GABVc80) | str. C6706, phage VP882 Δ*q*::$Cm^R$ | This study |
| *V. cholerae* (GABVc248) | str. C6706, phage VP882 Δ$traR_{VP882}$Δ*q*::$Cm^R$ | This study |
| *V. cholerae* (GABVc82) | str. C6706, phage VP882 Δ*qisA*Δ$traR_{VP882}$Δ*q*::$Cm^R$ | This study |
| *V. parahaemolyticus* 882 | Wild-type, phage VP882 lysogen | [4] |
| *E. coli* TOP10 | *F– mcrA Δ(mrr-hsdRMS-mcrBC) φ80lacZΔM15 ΔlacX74 recA1 araD139 Δ(ara-leu)7697 galU galK λ– rpsL(*$Str^R$*) endA1 nupG* | Invitrogen |
| *E. coli* BL21(DE3) | *B F- dcm ompT hsdS(rB- mB-) gal λ(DE3)* | Agilent |
| *E. coli* S17 | *ΔlacU169 (φlacZΔM15) recA1 endA1 hsdR17 thi-1 gyrA96 relA1 λpir* | [5] |
| *Strains carrying plasmids* | | |
| **Strain (ID)** | **Phage/Plasmid** | **Reference** |
| *V. cholerae* (GABVc104) | phage VP882 $Cm^R$ (WT phage VP882)/pTetV | This study |
| *V. cholerae* (GABVc102) | phage VP882 $Cm^R$ (WT phage VP882)/pGB121 | This study |
| *V. cholerae* (GABVc99) | phage VP882 $Cm^R$ (WT phage VP882)/pJES-143 | This study |
| *V. cholerae* (GABVc124) | phage VP882 Δ*q*::$Cm^R$/pJES-143 | This study |
| *V. cholerae* (GABVc122) | phage VP882 Δ*q*::$Cm^R$/pTetV | This study |
| *V. cholerae* (GABVc121) | phage VP882 Δ*q*::$Cm^R$/pGB121 | This study |
| *V. cholerae* (GABVc354) | pGB306 | This study |
| *V. cholerae* (GABVc238) | phage VP882 Δ*q*::$Cm^R$/pTetV, pJES-105 | This study |
| *V. cholerae* (GABVc239) | phage VP882 Δ*q*::$Cm^R$/pJES-143, pJES-105 | This study |

| | | |
|---|---|---|
| *V. cholerae* (GABVc332) | phage VP882 $Cm^R$ (WT phage VP882)/pTetV, pJES-105 | This study |
| *V. cholerae* (GABVc331) | phage VP882 $Cm^R$ (WT phage VP882)/pJES-143, pJES-105 | This study |
| *V. cholerae* (GABVc206) | phage VP882 Δ*qisA*Δ*traR*$_{VP882}$Δ*q*::$Cm^R$/pTetV | This study |
| *V. cholerae* (GABVc207) | phage VP882 Δ*qisA*Δ*traR*$_{VP882}$Δ*q*::$Cm^R$/pJES-143 | This study |
| *V. cholerae* (GABVc255) | phage VP882 Δ*traR*$_{VP882}$Δ*q*::$Cm^R$/pJES-143 | This study |
| *V. cholerae* (GABVc257) | phage VP882 Δ*traR*$_{VP882}$Δ*q*::$Cm^R$/pTetV | This study |
| *V. cholerae* (GABVc176) | phage VP882 $Cm^R$ (WT phage VP882)/pJES-052 | This study |
| *V. cholerae* (GABVc265) | phage VP882 $Cm^R$ (WT phage VP882)/pJY-014 | This study |
| *V. cholerae* (GABVc263) | phage VP882 Δ*q*::$Cm^R$/pJY-014 | This study |
| *V. cholerae* (GABVc264) | phage VP882 Δ*qisA*Δ*traR*$_{VP882}$Δ*q*::$Cm^R$/pJY-014 | This study |
| *V. cholerae* (GABVc266) | phage VP882 *qtiQ*::Tn*5*/pJY-014 | This study |
| *V. cholerae* (GABVc267) | phage VP882 Δ*qtip*::$Cm^R$/pJY-014 | This study |
| *V. cholerae* (GABVc268) | phage VP882 *q*::Tn*5*/pJY-014 | This study |
| *V. cholerae* (GABVc373) | phage VP882 Δ*traR*$_{VP882}$Δ*q*::$Cm^R$/pJY-014 | This study |
| *V. cholerae* (GABVc374) | phage VP882 Δ*traR*$_{VP882}$Δ*q*::$Cm^R$/pJES-052 | This study |
| *V. cholerae* (GABVc175) | phage VP882 Δ*qisA*Δ*traR*$_{VP882}$Δ*q*::$Cm^R$/pJES-052 | This study |
| *V. cholerae* (GABVc177) | phage VP882 *qtiQ*::Tn*5*/pJES-052 | This study |
| *V. cholerae* (GABVc178) | phage VP882 Δ*qtip*::$Cm^R$/pJES-052 | This study |
| *V. cholerae* (GABVc180) | phage VP882 *q*::Tn*5*/pJES-052 | This study |
| *V. cholerae* (GABVc183) | phage VP882 Δ*q*::$Cm^R$/pJES-052 | This study |
| *V. cholerae* (GABVc160) | phage VP882 *q*::Tn*5*/pJES-143 | This study |
| *V. cholerae* (GABVc164) | phage VP882 *q*::Tn*5*/pTetV | This study |
| *V. cholerae* (GABVc200) | phage VP882 $Cm^R$ (WT phage VP882)/pJES-052, pJES-105 | This study |
| *V. cholerae* (GABVc204) | phage VP882 *q*::Tn*5*/pJES-052, pJES-105 | This study |

| | | |
|---|---|---|
| *V. cholerae* (GABVc205) | phage VP882 Δ*q*::Cm$^R$/pJES-052, pJES-105 | This study |
| *V. cholerae* (GABVc339) | phage VP882 Δ*q*::Cm$^R$/pJY-014, pJES-105 | This study |
| *V. cholerae* (GABVc343) | phage VP882 Cm$^R$ (WT phage VP882)/pJY-014, pJES-105 | This study |
| *V. cholerae* (GABVc344) | phage VP882 *qtiQ*::Tn*5*/pJY-014, pJES-105 | This study |
| *V. cholerae* (GABVc345) | phage VP882 Δ*qtip*::Cm$^R$/pJY-014, pJES-105 | This study |
| *V. cholerae* (GABVc346) | phage VP882 *q*::Tn*5*/pJY-014, pJES-105 | This study |
| *V. cholerae* (GABVc492) | phage VP882 *qtiQ*::Tn*5*/pJES-052, pJES-105 | This study |
| *V. cholerae* (GABVc202) | phage VP882 Δ*qtip*::Cm$^R$/pJES-052, pJES-105 | This study |
| *V. cholerae* (GABVc358) | phage VP882 Δ*q*::Cm$^R$/pJES-052, pGB328 | This study |
| *V. cholerae* (GABVc359) | phage VP882 Δ*q*::Cm$^R$/pJY-014, pGB328 | This study |
| *V. cholerae* (GABVc381) | phage VP882 Δ*q*::Cm$^R$/pJES-052, pGB313 | This study |
| *V. cholerae* (GABVc382) | phage VP882 Δ*q*::Cm$^R$/pJY-014, pGB313 | This study |
| *V. cholerae* (GABVc393) | phage VP882 Δ*q*::Cm$^R$/pJES-052, pGB351 | This study |
| *V. cholerae* (GABVc394) | phage VP882 Δ*q*::Cm$^R$/pJY-014, pGB351 | This study |
| *V. cholerae* (GABVc399) | phage VP882 Δ*q*::Cm$^R$/pJES-052, pGB352 | This study |
| *V. cholerae* (GABVc400) | phage VP882 Δ*q*::Cm$^R$/pJY-014, pGB352 | This study |
| *V. cholerae* (GABVc503) | phage VP882 Δ*q*::Cm$^R$/pJES-052, pGB495 | This study |
| *V. cholerae* (GABVc504) | phage VP882 Δ*q*::Cm$^R$/pJY-014, pGB495 | This study |
| *V. choleraeΔtdh* (GABVc464) | pJY-014, pGB388 | This study |
| *V. choleraeΔtdh* (GABVc467) | pJES-052, pGB388 | This study |
| *V. cholerae* (GABVc468) | phage VP882 Δ*q*::Cm$^R$/pGB391 | This study |
| *V. cholerae* (GABVc470) | phage VP882 Δ*q*::Cm$^R$/pGB474 | This study |
| *V. cholerae* (GABVc471) | phage VP882 Cm$^R$ (WT phage VP882)/pGB391 | This study |
| *V. cholerae* (GABVc473) | phage VP882 Cm$^R$ (WT phage VP882)/pGB474 | This study |
| *V. cholerae* (GABVc486) | phage VP882 Δ*q*::Cm$^R$/pGB390 | This study |

| | | |
|---|---|---|
| *V. cholerae* (GABVc487) | phage VP882 $Cm^R$ (WT phage VP882)/pGB390 | This study |
| *V. parahaemolyticus* 882 | Wild-type, phage VP882 lysogen/pMMB*sacBtfoX* | This study |
| *E. coli* TOP10 (GABEc1096) | pGB388, pJES-052 | This study |
| *E. coli* TOP10 (GABEc1097) | pGB388, pJY-014 | This study |
| *E. coli* TOP10 (GABEc1098) | pGB388, pKP-375 | This study |
| *E. coli* BL21(DE3) (GABEc914) | pGB378 | This study |
| *E. coli* BL21(DE3) (GABEc917) | pGB381 | This study |
| *E. coli* BL21(DE3) (GABEc1431) | pGB519 | This study |
| *E. coli* BL21(DE3) (GABEc1432) | pGB519 | This study |
| *E. coli* BL21(DE3) (GABEc1433) | pGB519 | This study |
| *E. coli* BL21(DE3) (GABEc1434) | pGB520 | This study |
| *E. coli* BL21(DE3) (GABEc1435) | pGB520 | This study |
| *E. coli* BL21(DE3) (GABEc1436) | pGB520 | This study |
| *E. coli* BL21(DE3) (GABEc1440) | pGB378 | This study |
| *E. coli* BL21(DE3) (GABEc1441) | pGB378 | This study |
| *E. coli* BL21(DE3) (GABEc1442) | pGB381 | This study |
| *E. coli* BL21(DE3) (GABEc1443) | pGB381 | This study |
| *E. coli* TOP10 (JSS-1478) | pXB-300 | [2] |
| *E. coli* TOP10 (GABEc904) | pGB362 | This study |
| *E. coli* TOP10 (GABEc905) | pGB365 | This study |
| *E. coli* TOP10 (GABEc906) | pGB366 | This study |
| *E. coli* TOP10 (GABEc927) | pGB402 | This study |
| *E. coli* TOP10 (GABEc929) | pGB416 | This study |
| *E. coli* TOP10 (GABEc930) | pGB417 | This study |
| *E. coli* TOP10 (GABEc931) | pGB418 | This study |
| *E. coli* TOP10 (GABEc998) | pGB459 | This study |
| *E. coli* TOP10 (GABEc1044) | pGB476 | This study |
| *E. coli* TOP10 (GABEc1079) | pGB475 | This study |
| *E. coli* TOP10 (GABEc1082) | pGB364 | This study |
| *E. coli* TOP10 (GABEc1104) | pGB507 | This study |
| *E. coli* TOP10 (GABEc1104) | pGB503 | This study |
| *E. coli* TOP10 (GABEc1225) | pGB502 | This study |
| *E. coli* TOP10 (GABEc1248) | pGB510 | This study |
| *E. coli* TOP10 (GABEc1396) | pGB389, pGB416 | This study |
| *E. coli* TOP10 (GABEc1397) | pJY-014, pGB416 | This study |

| | | |
|---|---|---|
| *E. coli* TOP10 (GABEc1398) | pGB389, pGB417 | This study |
| *E. coli* TOP10 (GABEc1399) | pJY-014, pGB417 | This study |
| *E. coli* TOP10 (GABEc1400) | pGB389, pGB362 | This study |
| *E. coli* TOP10 (GABEc1401) | pJY-014, pGB362 | This study |
| *E. coli* TOP10 (GABEc1426) | pGB389, pXB-300 | This study |
| *E. coli* S17 | pRK2013 | Bassler lab Collection |

### Supplementary Table 2: Primers and gene blocks used in this study

| ID | Sequence (5' → 3') | Purpose |
|---|---|---|
| *Primers for recombinant phage generation* | | |
| ODO_493 | TGACAAGCAAGTCACTCAGAAATACATC | Colony PCR to verify VP882Δ*q* (FWD) |
| oGAB009 | TTCGAGGATAGCAGAGTCGTCCGC | Amplify 3kb up of phage VP882 neutral locus for PCR construct to generate VP882 $Cm^R$ (WT phage VP882) (FWD) |
| oGAB010 | AATCCTGACTGAGTTCGACC | SOE to generate VP882 $Cm^R$ (WT phage VP882) (FWD) |
| oGAB011 | TTGCACGCCGCGCTAACGCCGCGGCG CCGCTAGTTACGCCCCGCCCTGCCACT CATCGC | Amplify $Cm^R$, oriT for PCR construct to generate VP882 $Cm^R$ (WT phage VP882) (FWD) |
| oGAB012 | GCGATGAGTGGCAGGGCGGGGCGTAA CTAGCGGCGCCGCGGCGTTAGCGCGG CGTGCAA | Amplify 3kb up of phage VP882 neutral locus for PCR construct to generate VP882 $Cm^R$ (WT phage VP882) (REV) |
| oGAB013 | CCGCCCGAGGGCAGAGCCATGATTTTC TCCACTTCTAGCGTCTGTGCCCATCATT CGGGC | Amplify 3kb down of phage VP882 neutral locus for PCR construct to generate VP882 $Cm^R$ (WT phage VP882) (FWD) |
| oGAB014 | GGCACAGACGCTAGAAGTGGAGAAAAT CATGGCTCTGCCCTCGGGCGGACCAC GCCCA | Amplify $Cm^R$, oriT for PCR construct to generate VP882 $Cm^R$ (WT phage VP882) (REV) |
| oGAB015 | GCAATGACCAGCCGTCCTCGGCCG | Amplify 3kb down of phage VP882 neutral locus for PCR construct to generate VP882 $Cm^R$ (WT phage VP882) (REV) |
| oGAB016 | GTGCGGTGGTTGAACTCTTC | SOE to generate VP882 $Cm^R$ (WT phage VP882) (REV) |
| oGAB028 | CGCAGGCGCTGCAGGCAGTAC | Amplify 3kb up of phage VP882 *q* for PCR construct to generate VP882Δ*q* (FWD) |
| oGAB029 | GACGACCTTTTTGAGGTTG | SOE to generate VP882Δ*q* (FWD) |
| oGAB032 | GGGCGTGGTCCGCCCGAGGGCAGAG CCATGACGCTATGAACTGCACACACTG CGGCACTG | Amplify 3kb down of phage VP882 *q* for PCR construct to generate recombinant phages missing *q* (FWD) |

| | | |
|---|---|---|
| oGAB033 | AGTGCCGCAGTGTGTGCAGTTCATAGCGTCATGGCTCTGCCCTCGGGCGGACCACGCCC | Amplify $Cm^R$, oriT for PCR construct to generate recombinant phages missing *q* (REV) |
| oGAB034 | AGTGATACCCATAAGCAAG | Amplify 3kb down of phage VP882 *q* for PCR construct to generate recombinant phages missing *q* (REV) |
| oGAB035 | TCAATGCGTGAAGTGTTG | SOE to generate recombinant phages missing *q* (REV) |
| oGAB104 | AGGCGCTTATAGTGGTACACGTG | Colony PCR to verify recombinant phages missing *q* (REV) |
| oGAB199 | AGGTATGGAAGAAGGGACTCG | Colony PCR to verify VP882 $Cm^R$ (WT phage VP882) (FWD) |
| oGAB200 | CGTCTATTACTTCGCTATCGGATG | Colony PCR to verify VP882 $Cm^R$ (WT phage VP882) (REV) |
| oGAB233 | TCTGCAGGAGATTAAAGGCCGACATCAAAGGTAGTTACGCCCCGCCCTGCCACTCATCGC | Amplify $Cm^R$, oriT for PCR construct to generate VP882Δ*q* (FWD) |
| oGAB234 | ACTGCGATGAGTGGCAGGGCGGGGCGTAACTACCTTTGATGTCGGCCTTTAATCTCCTGC | Amplify 3kb up of phage VP882 *q* for PCR construct to generate VP882Δ*q* (REV) |
| oGAB237 | GCGTCACACATAACGACGCGGCGGG | Amplify 3kb up of phage VP882 *qisA-traR*$_{VP882}$-*q* for PCR construct to generate VP882 Δ*qisA*Δ*traR*$_{VP882}$Δ*q* (FWD) |
| oGAB238 | TCTCCTGGTAGTTAGCAATGCC | SOE to generate VP882 Δ*qisA*Δ*traR*$_{VP882}$Δ*q* (FWD) |
| oGAB241 | CAATGTTGCGTGCTATGTAAACGG | Colony PCR to verify VP882 Δ*qisA*Δ*traR*$_{VP882}$Δ*q* (FWD) |
| oGAB248 | AGCGTTGTGCAGTTTCCCTTTGG | Amplify 3kb up of phage VP882 *traR*$_{VP882}$-*q* for PCR construct to generate VP882 Δ*traR*$_{VP882}$Δ*q* (FWD) |
| oGAB249 | GCTGCTCTTCATCCAGTTCGC | SOE to generate VP882 Δ*traR*$_{VP882}$Δ*q* (FWD) |
| oGAB250 | TCTCGATGTTCCGCTACC | Colony PCR to verify VP882 Δ*traR*$_{VP882}$Δ*q* (FWD) |
| oGAB253 | GGTGCCCAACACTTAGTAATGGAAACCCGGAAAATTACGCCCCGCCCTGCCACTCATCGC | Amplify $Cm^R$, oriT for PCR construct to generate VP882 Δ*qisA*Δ*traR*$_{VP882}$Δ*q* (FWD) |
| oGAB254 | AACAGTACTGCGATGAGTGGCAGGGCGGGGCGTAATTTTCCGGGTTTCCATTACTAAGTG | Amplify 3kb up of phage VP882 *qisA-traR*$_{VP882}$-*q* for PCR construct to generate VP882 Δ*qisA*Δ*traR*$_{VP882}$Δ*q* (REV) |
| oGAB264 | TGTCAGTTCAACTAACGGAAGAAACCCAATGATTACGCCCCGCCCTGCCACTCATCGCAG | Amplify CmR, oriT for PCR construct to generate VP882 Δ*traR*$_{VP882}$Δ*q* (FWD) |
| oGAB265 | ACTGCGATGAGTGGCAGGGCGGGGCGTAATCATTGGGTTTCTTCCGTTAGTTGAACTGAC | Amplify 3kb up of phage VP882 *traR*$_{VP882}$-*q* for PCR construct to |

| | | |
|---|---|---|
| | | generate VP882 $\Delta traR_{VP882}\Delta q$ (REV) |
| *Primers for plasmid assembly* | | |
| oGAB906 | GGAGAAAGGGGCTACCGGCGATGATCGGCACGTAAGAGG | Amplify insert ($Cm^R$) for pGB306 (FWD) |
| oGAB909 | CTTTATAAAGGGGCTGCTGGTTTACGCCCCGCCCTGCC | Amplify insert ($Cm^R$) for pGB306 (REV) |
| oGAB907 | CCTCTTACGTGCCGATCATCGCCGGTAGCCCCTTTC | Amplify backbone (pJES-143) for pGB306 (FWD) |
| oGAB908 | GGCAGGGCGGGGCGTAAACCAGCAGCCCCTTTATAAAG | Amplify backbone (pJES-143) for pGB306 (REV) |
| oGAB490 | CTCTTTAGCCGCCCTACAATCCAGCCTG | Introduce point mutant (*qtip D13A*) to generate pGB121 (FWD) |
| oGAB491 | AGGGCGGCTAAAGAGTCGATGTGTTCACTGC | Introduce point mutant (*qtip D13A*) to generate pGB121 (REV) |
| oGAB888 | GCTTGCCTCGGTTCTTGACCTAGAGGTCCCCTTTTTTATTTAAAAATTTTTTCAC | Amplify backbone (pJES-105) for pGB313, pGB328 (FWD) |
| oGAB891 | GGAACAAAAGGACGAATGACGTCCCCAATTACTAGCCAATTCG | Amplify backbone (pJES-105) for pGB313, pGB328 (REV) |
| oGAB889 | GGGGACCTCTAGGTCAAGAACCGAGGCAAGC | Amplify MRR insert for pGB313, pGB328 (FWD) |
| oGAB890 | GGCTAGTAATTGGGGACGTCATTCGTCCTTTTGTTCCTTC | Amplify MRR insert for pGB313, pGB328 (REV) |
| oGAB947 | GATTTGCTAGCATTCGTCTCCAGCGCG | Generate fragment from pGB313 (1 of 2) for pGB351 (FWD) |
| oGAB953 | TATCATTGCAGCACTGGGG | Generate fragment from pGB313 (1 of 2) for pGB351, pGB352, pGB495 (REV) |
| oGAB948 | GAATGCTAGCAAATCCTGACTGAGTTCGACC | Generate fragment from pGB313 (2 of 2) for pGB351 (FWD) |
| oGAB954 | CGATACGGGAGGGCTTACC | Generate fragment from pGB313 (2 of 2) for pGB351, pGB352, pGB495 (REV) |
| oGAB949 | GCCTTGTGATGCCTGGCATACATCGAACAAGG | Generate fragment from pGB313 (2 of 2) for pGB352 (FWD) |
| oGAB950 | CAGGCATCACAAGGCGCCCTGGGCG | Generate fragment from pGB313 (1 of 2) for pGB352 (FWD) |
| oGAB900 | GTTAGGCAACGTGTAGGTCACAAATCACC | Generate fragment (1 of 2) from VP882 $Cm^R$ plasmid for insert template for pGB328 (FWD) |
| oGAB764 | CATCAGCACCTTGTCGCC | Generate fragment (1 of 2) from VP882 $Cm^R$ plasmid for insert template for pGB328 (REV) |
| oGAB901 | GGTGATTTGTGACCTACACGTTGCCTAAC | Generate fragment (2 of 2) from VP882 $Cm^R$ plasmid for insert template for pGB328 (FWD) |
| oGAB767 | GGCAAATATTATACGCAAGGCG | Generate fragment (2 of 2) from VP882 $Cm^R$ plasmid for insert template for pGB328 (REV) |

| | | |
|---|---|---|
| oGAB1047 | TCGATGTGGGCCAGGCACACAAGGCGC | Generate fragment from pGB313 (2 of 2) for pGB495 (FWD) |
| oGAB1048 | CCTGGCCCACATCGAACAAGGAAAACGACGC | Generate fragment from pGB313 (1 of 2) for pGB495 (FWD) |
| oGAB868 | CTAGTACAGGAGGTGTGAAGCTGTTCAACCAGCTCGTCATAACGG | Generate fragment from pJES-105 (1 of 2) for pGB388 (FWD) |
| oGAB1086 | CCGTTAATAATGAATGAAATTTTTTAGTCATCCTGGCATACATCGAACAAG | Generate fragment from pJES-105 (1 of 2) for pGB388 (REV) |
| oGAB869 | CCGTTATGACGAGCTGGTTGAACAGCTTCACACCTCCTGTACTAG | Generate fragment from pJES-105 (2 of 2) for pGB388 (FWD) |
| oGAB1085 | CTTGTTCGATGTATGCCAGGATGACTAAAAAAATTTCATTCATTATTAACGG | Generate fragment from pJES-105 (2 of 2) for pGB388 (REV) |
| oGAB1006 | CTTAAATGTGAAAGTGGGTCTTAATCCCGTCAAGTCAGCGTAATG | Amplify backbone for pGB390 (from pGB389), pGB391 (from pJY-014), pGB474 (from pGB389) (FWD) |
| oGAB1007 | CGCTCTCCTGAGTAGGACAAATCGGCGGCTTTGTTGAATAAATC | Amplify backbone for pGB390 (from pGB389), pGB391 (from pJY-014), pGB474 (from pGB389) (REV) |
| oGAB1005 | CATTACGCTGACTTGACGGGATTAAGACCCACTTTCACATTTAAG | Amplify insert for pGB390 ($P_{tetA}$-*qtip*), pGB391 ($P_{tetA}$-*qtip*), pGB474 ($P_{tetA}$-empty) (FWD) |
| oGAB1008 | GATTTATTCAACAAAGCCGCCGATTTGTCCTACTCAGGAGAGCG | Amplify insert for pGB390 ($P_{tetA}$-*qtip*), pGB391 ($P_{tetA}$-*qtip*), pGB474 ($P_{tetA}$-empty) (REV) |
| oGAB1000 | GGCTGTTCTGGTGTTGCTAGGGCTGCTGCCACCGCTG | Amplify backbone (pGB375) for pGB378 (FWD) |
| oGAB1001 | CCCGAAAAGTGCCACCTGTAGGTTAATTAAGCTCAGCGGTTGAC | Amplify backbone (pGB375) for pGB378 (REV) |
| oGAB999 | CAGCGGTGGCAGCAGCCCTAGCAACACCAGAACAGCC | Amplify insert ($HIS_6$-$TraR_{VP882}$) for pGB378 (FWD) |
| oGAB1002 | GCTTAATTAACCTACAGGTGGCACTTTTCGGG | Amplify insert ($HIS_6$-$TraR_{VP882}$) for pGB378 (REV) |
| oGAB996 | GTTCAACTAACGGAAGAAACCCAAACCACTGAGGATCTGTACTTTCAG | Amplify backbone (pRSF1b) for pGB375 (FWD) |
| oGAB998 | GATAATTATTTGGTCTTCGGTCATGGTATATCTCCTTATTAAAGTTAAACAAAATTATTT | Amplify backbone (pRSF1b) for pGB375 (REV) |
| oGAB995 | CTGAAAGTACAGATCCTCAGTGGTTTGGGTTTCTTCCGTTAGTTGAAC | Amplify insert (*qisA*) for pGB375 (FWD) |
| oGAB997 | AAATAATTTTGTTTAACTTTAATAAGGAGATATACCATGACCGAAGACCAAATAATTATC | Amplify insert (*qisA*) for pGB375 (REV) |
| oGAB993 | ATACCATGACCACTGAGGATCTGTACTTT | Generate pGB381 from pGB378 (FWD) |
| oGAB994 | CAGTGGTCATGGTATATCTCCTTATTAAAGTT | Generate pGB381 from pGB378 (REV) |
| oGAB1163 | GGTATGAGTAATATTGCAAATCAGGCACAGGACGTCATCG | Generate pGB519 (from pGB378), pGB520 (from pGB381) (FWD) |

| | | |
|---|---|---|
| oGAB1164 | CGTCCTGTGCCTGATTTGCAATATTACT CATACCGCTGCTACCG | Generate pGB519 (from pGB378), pGB520 (from pGB381) (REV) |
| oGAB1024 | GGCTTGCCTCGGTTCTTGACGCAACCA GGATCCGGTG | Amplify backbone (pJY-014) for pGB389 (FWD) |
| oGAB1025 | GCCTTGTGTGCCTGGCATTTCACACCT CCTGCAGGTACC | Amplify backbone (pJY-014) for pGB389 (REV) |
| oGAB1023 | CACCGGATCCTGGTTGCGTCAAGAACC GAGGCAAGCC | Amplify insert (*qtiQ*) for pGB389 (FWD) |
| oGAB1026 | GGTACCTGCAGGAGGTGTGAAATGCCA GGCACACAAGGC | Amplify insert (*qtiQ*) for pGB389 (REV) |
| oGAB964 | CGATAATTATTTGGTCTTCGGTCATTTCA CACCTCCTGTGGAG | Amplify backbone (pXB-300) for pGB362, pGB418 (REV) |
| oGAB965 | GGCCGACATCAAAGGTAGACCCTCAGC CAATGCGC | Amplify backbone (pXB-300) for pGB362 (FWD) |
| oGAB963 | CTCCACAGGAGGTGTGAAATGACCGAA GACCAAATAATTATCG | Amplify insert for pGB362 (*qisA*-*traR*$_{VP882}$), pGB418 (*qisA*-*traR*$_{VP882}$-*q*), (REV) |
| oGAB966 | GCGCATTGGCTGAGGGTCTACCTTTGA TGTCGGCC | Amplify insert (*qisA*-*traR*$_{VP882}$) for pGB362 (FWD) |
| oGAB971 | GTGACCGACACGTTGCCTAAACCCTCA GCCAATGCGCTG | Amplify backbone (pXB-300) for pGB365, pGB366, pGB418, (FWD) |
| oGAB974 | GTGCCTGATCTGCAATATCACTCATTTC ACACCTCCTGTGGAGCTCC | Amplify backbone (pXB-300) for pGB365 (REV) |
| oGAB972 | CAGCGCATTGGCTGAGGGTTTAGGCAA CGTGTCGGTCAC | Amplify insert for pGB365 (*traR*$_{VP882}$-*q*), pGB418 (*qisA*-*traR*$_{VP882}$-*q*) (FWD) |
| oGAB973 | GGAGCTCCACAGGAGGTGTGAAATGA GTGATATTGCAGATCAGGCAC | Amplify insert for pGB365 (*traR*$_{VP882}$-*q*), pGB366 (*q*) (REV) |
| oGAB976 | GCTCGTTGTATTCCCGTACTCTCATTTC ACACCTCCTGTGGAGCTCC | Amplify backbone (pXB-300) for pGB366 (REV) |
| oGAB975 | GGAGCTCCACAGGAGGTGTGAAATGA GAGTACGGGAATACAACGAGC | Amplify insert for pGB366 (*q*) (FWD) |
| oGAB961 | CCCAATGAATGAGAGTACGGGAATACA ACGAGC | Generate pGB364 (*qisA*-*traR*$_{VP882}$-*q* → *qisA, q*) (FWD) |
| oGAB962 | CTCTCATTCATTGGGTTTCTTCCGTTAG TTG | Generate pGB364 (*qisA*-*traR*$_{VP882}$-*q* → *qisA, q*) (REV) |
| oGAB1039 | GTAATATTGCAAATCAGGCACAGGACGT CATCG | Introduce NxxNxA for pGB402, pGB502, pGB503 (FWD) |
| oGAB1040 | GTCCTGTGCCTGATTTGCAATATTACTC ATTGGGTTTCTTCCGTTAG | Introduce NxxNxA for pGB402, pGB502, pGB503 (REV) |
| oGAB1053 | CCCAATGAACCCTCAGCCAATGCGCT | Generate pGB416 from pGB362 (FWD) |
| oGAB1054 | TGAGGGTTCATTGGGTTTCTTCCGTTA GTTGA | Generate pGB416 from pGB362 (REV) |
| oGAB1055 | GGTGTGAAATGAGTGATATTGCAGATCA GGCA | Generate pGB417 from pGB362 (FWD) |
| oGAB1056 | CACTCATTTCACACCTCCTGTGGAGCT | Generate pGB417 from pGB362 (REV) |

| | | |
|---|---|---|
| oGAB1110 | CTTCATCGGCTTCATCACTCATTTCACA CCTCCTGTGGAGC | Amplify backbone (pXB-300) for pGB475 (FWD) |
| oGAB1111 | GGAAAGACAGAGAAAACATTATGCATAA ACCCTCAGCCAATGCGCTG | Amplify backbone (pXB-300) for pGB475 (REV) |
| oGAB1109 | GCTCCACAGGAGGTGTGAAATGAGTGA TGAAGCCGATGAAG | Amplify insert (*traR*$_F$) for pGB475 (FWD) |
| oGAB1112 | CAGCGCATTGGCTGAGGGTTTATGCAT AATGTTTTCTCTGTCTTTCC | Amplify insert (*traR*$_F$) for pGB475 (REV) |
| oGAB1114 | GCTGAATCAATGATGTCTGCCATTTCAC ACCTCCTGTGGAGC | Amplify backbone (pXB-300) for pGB476 (FWD) |
| oGAB1115 | CAGTAAACAGAGAGGTTCGAAGTAAAC CCTCAGCCAATGCGCTG | Amplify backbone (pXB-300) for pGB476 (REV) |
| oGAB1113 | GCTCCACAGGAGGTGTGAAATGGCAG ACATCATTGATTCAGC | Amplify insert (*traR*$_\lambda$) for pGB476 (FWD) |
| oGAB1116 | CAGCGCATTGGCTGAGGGTTTACTTCG AACCTCTCTGTTTACTG | Amplify insert (*traR*$_\lambda$) for pGB476 (REV) |
| oGAB1067 | GTAATATTGCAAATCAGGCACAGGACGT CATC | Generate pGB459 from pGB417 (FWD) |
| oGAB1068 | GATTTGCAATATTACTCATTTCACACCTC CTGT | Generate pGB459 from pGB417 (REV) |
| oGAB1130 | CGGTGTAGCTCTTTGGCATCATTGGGT TTCTTCCGTTAG | Amplify backbone (pGB416) for pGB507 (FWD) |
| oGAB1131 | GGAAGATCACTTCGCAGAATAAACCCT CAGCCAATGCGC | Amplify backbone for pGB507 (pGB416), pGB510 (pXB-300) (REV) |
| oGAB1129 | CTAACGGAAGAAACCCAATGATGCCAA AGAGCTACACCG | Amplify insert (*traK*) for pGB507 (FWD) |
| oGAB1132 | GCGCATTGGCTGAGGGTTTATTCTGCG AAGTGATCTTCC | Amplify insert (*traK*) for pGB507, pGB510 (REV) |
| oGAB1147 | GCTCCACAGGAGGTGTGAAAATGCCAA AGAGCTACACCG | Amplify insert (*traK*) for pGB510 (FWD) |
| oGAB1148 | CGGTGTAGCTCTTTGGCATTTTCACAC CTCCTGTGGAGC | Amplify backbone (pXB-300) for pGB510 (FWD) |
| *Primers for RT-qPCR* | | |
| FJS-q009 | CTGCTGACTCTGATTGTGCTG | *gp69* (FWD) (courtesy of F. Santoriello) |
| FJS-q010 | TCGTGAGAGGTGATGTACTTCTC | *gp69* (REV) (courtesy of F. Santoriello) |
| oGAB669 | CTAAGGGGCAATCTCTACAAGACCCA | *V. cholerae hfq* (FWD) |
| oGAB670 | ACCATTTGGTTTACTGTGTTCTTCAGCA AG | *V. cholerae hfq* (REV) |
| *Gene blocks* | | |
| *traR*$_F$ | GGATCCTACCTGACGCTTTTTATCGCAA CTCTCTACTGTTTCTCCATACCCGTTTT TTGGGCTAACAGGAGGAATTCACCATG AGTGATGAAGCCGATGAAGCATATTCA GTGACAGAACAACTGACCATGACAGGA ATAAACCGGATACGCCAGAAAATAAATG CTCATGGTATTCCTGTTTATCTCTGTGA AGCATGCGGAAATCCTATTCCGGAAGC | Template for insert for pGB475 |

| | | |
|---|---|---|
| | CCGGCGGAAAATATTTCCCGGTGTGAC<br>GTTGTGCGTTGAATGTCAGGCGTATCA<br>GGAAAGACAGAGAAAACATTATGCATAA<br>AGCTTGGCTGTTTTGGCGGATGAGAG | |
| *traR*$_{\lambda}$ | GGATCCTACCTGACGCTTTTTATCGCAA<br>CTCTCTACTGTTTCTCCATACCCGTTTT<br>TTGGGCTAACAGGAGGAATTCACCATG<br>GCAGACATCATTGATTCAGCATCAGAAA<br>TAGAAGAATTACAGCGCAACACAGCAAT<br>AAAAATGCGCCGCCTGAACCACCAGGC<br>TATATCTGCCACTCATTGTTGTGAGTGT<br>GGCGATCCGATAGATGAACGAAGACGC<br>CTGGTCGTTCAGGGTTGTCGGACTTGT<br>GCAAGTTGCCAGGAGGATCTGGAACTT<br>ATCAGTAAACAGAGAGGTTCGAAGTAA<br>AGCTTGGCTGTTTTGGCGGATGAGAGA<br>AGATTTTCAGCCTGATACAGATTAAATC<br>AGAACGCAGAAGCGGTCTGATAAAA | Template for insert for pGB476 |
| $HIS_6$-<br>$TraR_{VP882}$ | CAGGTGGCACTTTTCGGGTAATACGAC<br>TCACTATAGGGGAATTGTGAGCGGATAA<br>CAATTCCCCTGTAGAAATAATTTTGTTTA<br>ACTTTAATAAGGAGATATACCATGCACC<br>ACCACCATCACCATCATCACCACCACG<br>GTAGCAGCGGTATGAGTGATATTGCAG<br>ATCAGGCACAGGACGTCATCGAGCAGC<br>ACCTGACGGCCAGCCTGGCGAACAGG<br>AAGCACAACATCAACCCAGCCATCCCA<br>AGCGCGAAGCATTGCGATGACTGCGA<br>GTCGGAAATCCCAGAGGCTCGCCGTC<br>GCAGTCTTCCTGGTGTCCGCTTGTGTG<br>TTGATTGCGCTTCTCTGCAGGAGATTAA<br>AGGCCGACATCAAAGGTAGGGCTGTTC<br>TGGTGTTGCTAG | Template for insert for pGB378 |

### Supplementary Table 3: Plasmids used in this study

| Plasmid name (informal) | Plasmid ID | Relevant Figures | Selection | Reference |
|---|---|---|---|---|
| pRK2013 | | (Methods) – Mating helper for conjugations | Kan | [6] |
| pMMB*sacBtfoX* | | (Methods) – Generation of recombinant phage | Kan | [7] |
| pRE112 | | (Methods) – PCR template for $Cm^R$, oriT for generating recombinant phage | Cm | [8] |

| | | | | |
|---|---|---|---|---|
| $P_{tetA}$-*qtip*, $Kan^R$ | pJES-143 | 2, Ext. Data 1, 2b | Kan | [2] |
| $P_{tetA}$-*qtip*, $Kan^R$, $Cm^R$ | pGB306 | 2a, Ext. Data 1a | Kan, Cm | This study |
| $P_{tetA}$-empty vector, $Kan^R$ | pTetV | 2b-e, Ext. Data 1, 2b | Kan | This study; courtesy of J. Valastyan |
| $P_{tetA}$-*qtip D13A*, $Kan^R$ | pGB121 | 2b, Ext. Data 1a | Kan | This study |
| $P_{BAD}$-$vqmA_{Phage}$ | pJES-052 | 3a, 3c, 3d, 5, Ext. Data 2a, 2c-e, 4a | Kan | [2] |
| $P_{BAD}$- empty vector | pJY-014 | 3a, 3c, 3d, 4c, 5, Ext. Data 2d-e, 4a-d | Kan | [2] |
| $P_{BAD}$-$vqmA_{Vc}$ | pKP-375 | 3d | Kan | [2] |
| P*gp69-lux* | pJES-105 | 2c, 3c, 5b, Ext. Data 1b, 2c-d, 4d | Crb | [2] |
| P*gp69-lux* + MRR | pGB313 | 3c, Ext. Data 2c-d | Crb | This study |
| P*gp69-lux* + MRR FS1 | pGB351 | 3c, Ext. Data 2c-d | Crb | This study |
| P*gp69-lux* + MRR FS2 | pGB352 | 3c, Ext. Data 2c-d | Crb | This study |
| P*gp69-lux* + MRR $\Delta gp64^{STOP}$ | pGB328 | 3c, Ext. Data 2c-d | Crb | This study |
| P*gp69-lux* + MRR $\Delta gp63^{ATG\rightarrow GGG}$ | pGB495 | 3c, Ext. Data 2c-d | Crb | This study |
| P*gp63-lux* | pGB388 | 3d, Ext. Data 2e | Crb | This study |
| $P_{BAD}$-*qtiQ*, $P_{tetA}$-*qtip* | pGB390 | 3e, Ext. Data 2f | Kan | This study |
| $P_{BAD}$-empty, $P_{tetA}$-*qtip* | pGB391 | 3e, Ext. Data 2f | Kan | This study |
| $P_{BAD}$-*qtiQ*, $P_{tetA}$-empty | pGB474 | 3e, Ext. Data 2f | Kan | This study |
| pRSF1b- $P_{T7}$-*qisA-HALO* | pGB375 | - | Kan | This study |
| pRSF1b- $P_{T7}$-*qisA-HALO*, $P_{T7}$-$HIS_6$-$traR_{VP882}$ | pGB378 | 4a | Kan | This study |
| pRSF1b- $P_{T7}$-*HALO*, $P_{T7}$-$HIS_6$-$traR_{VP882}$ | pGB381 | 4a | Kan | This study |
| pRSF1b- $P_{T7}$-*qisA-HALO*, $P_{T7}$-$HIS_6$-$traR^{NxxNxA}{}_{VP882}$ | pGB519 | 4a | Kan | This study |
| pRSF1b- $P_{T7}$-*HALO*, $P_{T7}$-$HIS_6$-$traR^{NxxNxA}{}_{VP882}$ | pGB520 | 4a | Kan | This study |
| $P_{tetA}$-empty vector, $Crb^R$ | pXB-300 | 4b-c, Ext. Data 3 | Crb | [9] |
| $P_{tetA}$-*qisA* | pGB416 | 4b-c, Ext. Data 3 | Crb | This study |
| $P_{tetA}$-$traR_{VP882}$ | pGB417 | 4b-c, Ext. Data 3 | Crb | This study |
| $P_{tetA}$-$traR^{NxxNxA}{}_{VP882}$ | pGB459 | 4b, Ext. Data 3a, 3c | Crb | This study |
| $P_{tetA}$-*qisA*-$traR_{VP882}$ | pGB362 | 4b-c, Ext. Data 3 | Crb | This study |
| $P_{tetA}$-*qisA*-$traR^{NxxNxA}{}_{VP882}$ | pGB402 | 4b, Ext. Data 3a, 3c | Crb | This study |
| $P_{tetA}$-$traR_F$ | pGB475 | Ext. Data 3b | Crb | This study |
| $P_{tetA}$-$traR_{\lambda}$ | pGB476 | Ext. Data 3b | Crb | This study |
| $P_{tetA}$-*traK* | pGB510 | Ext. Data 3b | Crb | This study |
| $P_{tetA}$-*qisA, traK* | pGB507 | Ext. Data 3b | Crb | This study |
| $P_{tetA}$-*q* | pGB366 | Ext. Data 3c | Crb | This study |
| $P_{tetA}$-$traR_{VP882}$-*q* | pGB365 | Ext. Data 3c | Crb | This study |
| $P_{tetA}$-$traR^{NxxNxA}{}_{VP882}$-*q* | pGB502 | Ext. Data 3c | Crb | This study |
| $P_{tetA}$-*qisA, q* | pGB364 | Ext. Data 3c | Crb | This study |
| $P_{tetA}$-*qisA*-$traR_{VP882}$-*q* | pGB418 | Ext. Data 3c | Crb | This study |

| | | | | |
|---|---|---|---|---|
| $P_{tetA}$-*qisA-traR*$^{NxxNxA}$$_{VP882}$-*q* | pGB503 | Ext. Data 3c | Crb | This study |
| $P_{BAD}$-*qtiQ* | pGB389 | 4c | Kan | This study |

### Supplementary Table 4: Time points for bar graphs

| Figure | Time (min) |
|---|---|
| 2b | 300 |
| 2c | 150 |
| 2d | 330 |
| 2e | 410 |
| 3a | 410 |
| 3c | 410 |
| 3d | 270 |
| 3e | 600 |
| 4b | 800 |
| 5b | 280 |
| 5c | 640 |
| Ext. Data 1e | Overnight (plated 200 min post induction) |
| Ext. Data 2d | 240 |
| Ext. Data 2e | 510 |
| Ext. Data 3d | 800 |
| Ext. Data 4a | 210 |
| Ext. Data 4c | 460 |
| Ext. Data 4d | 510 |
